# Precise Coordination of 3-dimensional Rotational Kinematics by Ventral Tegmental Area GABAergic Neurons

**DOI:** 10.1101/678391

**Authors:** Ryan N. Hughes, Glenn D.R. Watson, Elijah Petter, Namsoo Kim, Konstantin I. Bakhurin, Henry H. Yin

**Affiliations:** Department of Psychology and Neuroscience, Duke University, Durham, NC, 27708, USA; Department of Neurobiology, Duke University School of Medicine, Durham, NC, 27708, USA

## Abstract

The Ventral Tegmental Area (VTA) is a midbrain region implicated in a variety of motivated behaviors. However, the function of VTA GABAergic (Vgat+) neurons remains poorly understood. Here, using 3D motion capture, *in vivo* electrophysiology and calcium imaging, and optogenetics, we demonstrate a novel function of VTA^Vgat+^ neurons. We found three distinct populations of neurons, each representing head angle about a principal axis of rotation: pitch, roll, and yaw. For each axis, opponent cell groups were found that increase firing when the head moves in one direction, and decrease firing in the opposite direction. Selective excitation and inhibition of VTA^Vgat+^ neurons generate opposite rotational movements. The relationship between these neurons and head angle is degraded only at the time of reward consumption, at which point all head-angle related neuronal subpopulations show indistinguishable reward-related responses. Thus, VTA^Vgat+^ neurons serve a critical role in the control of rotational kinematics while pursuing a moving target. This general-purpose steering function can guide animals toward desired spatial targets in any motivated behavior.

## Introduction

The ventral tegmental area (VTA) has been implicated in motivated behaviors, addiction, and psychiatric disorders (Lammel et al., 2012; Volkow and Morales, 2015). Yet despite decades of research, the functional significance of the VTA is still poorly understood. Much research has focused on investigating the function of VTA dopamine neurons, especially in relation to behaviors guided by both reward and aversion (Eshel et al., 2015; Lammel et al., 2014; Tye et al., 2013). By comparison, fewer studies have investigated VTA gamma-aminobutyric acid (GABA) neurons, the other main cell type in the VTA, which comprises ~35% of its neural population (Fields et al., 2007; Nair-Roberts et al., 2008; Tan et al., 2012; van Zessen et al., 2012). Importantly, the less-studied GABAergic population contains many projection neurons, which can have significant inhibitory impact on their numerous downstream targets (Beier et al., 2019; Morales and Margolis, 2017; Taylor et al., 2014).

VTA GABAergic neurons are hypothesized to serve a role in aversive behaviors(Tan et al., 2012), as well as providing the subtraction necessary to compute prediction errors (Eshel et al., 2015). In addition, putative VTA GABAergic neurons have also been found to correlate with locomotion (Puryear et al., 2010; Wang and Tsien, 2011), and optogenetic activation of VTA GABA neurons disrupts reward-consumption(van Zessen et al., 2012). Clearly, VTA GABAergic neurons contribute to diverse behaviors, which is reflected in the heterogeneity of its anatomical connectivity. What is not clear, however, is how these reward- and locomotion-related processes interact and influence each other.

## Results

To investigate the role of VTA GABAergic neurons in behavior, we first wirelessly recorded single-unit activity in the VTA (*n* = 479) while mice performed a reward-tracking task during three-dimensional (3D) motion capture (Figure 1A) (Bartholomew et al., 2016; Fan et al., 2011). During this task, water-deprived mice were trained to continuously track a moving spout to receive a reward. The spout moved along either the horizontal (left and right) or vertical (up and down) axis during each session (Figure 1A). Kinematic data was extracted from markers placed on the head by infrared cameras, allowing quantification of behavioral variables such as position, velocity, acceleration, and head angle changes during continuous behavior.

**Figure 1.**
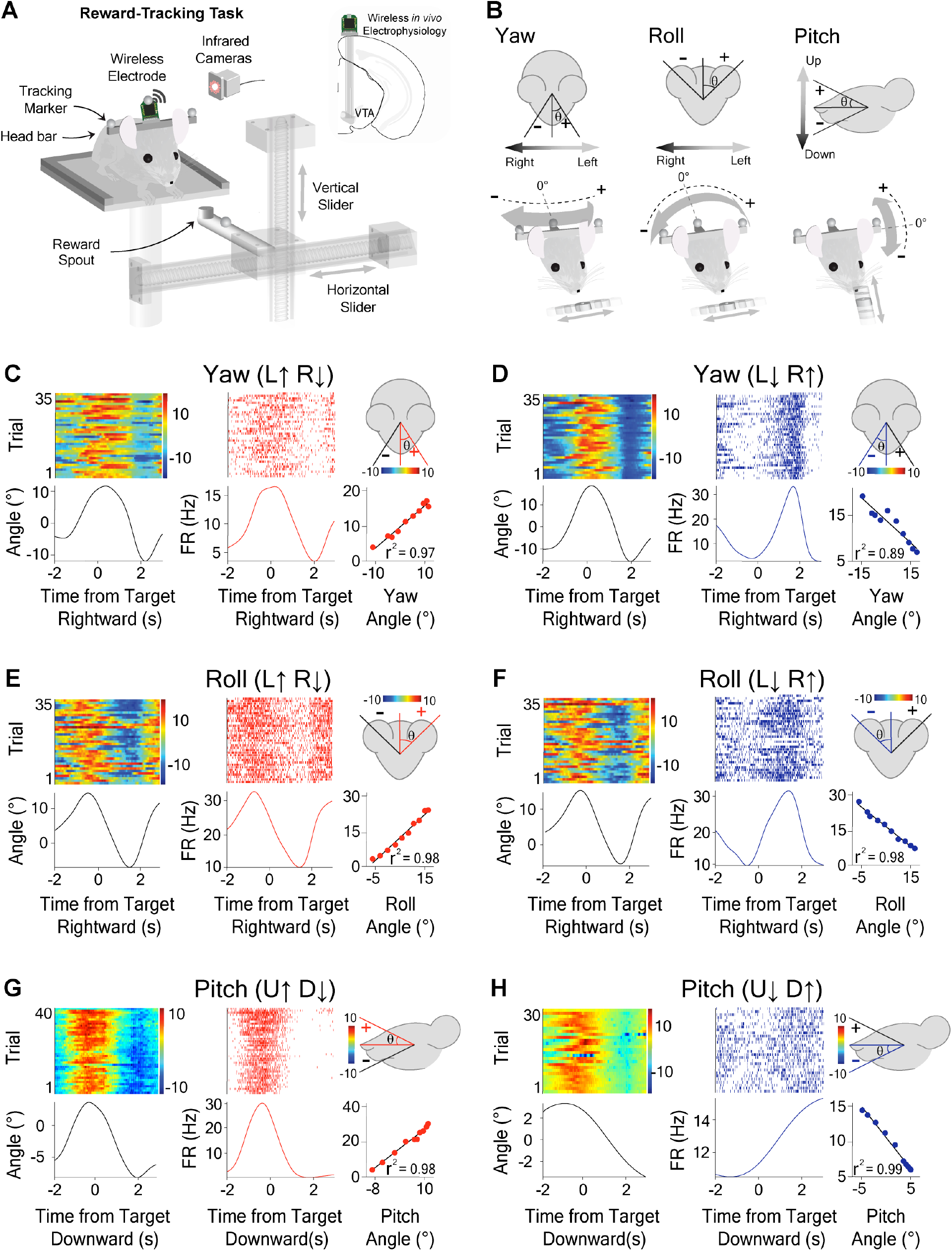
Opponent encoding of yaw, roll, and pitch angles by VTA GABAergic neurons. **(A)** Schematic of reward-tracking task during VTA wireless *in vivo* electrophysiology. Mice tracked a motorized spout in the horizontal and vertical directions to receive a reward (diluted condensed milk). Infrared cameras captured kinematic data from markers on the head of the animal with respect to reward target. Inset shows schematic of electrode placement into the VTA. **(B)** Schematic representation and sign conventions for yaw (*left*), roll (*middle*) and pitch (*right*) rotational kinematics. Arrows show direction of head movement around each axis. **(C-H)**Individual neurons represent direction specific angles along orthogonal axes of rotation. Peri-event heat maps of head angle (*left*), peri-event raster plots of VTA GABAergic neural activity (*middle)* with respect to reward target, and a correlation graph (*right bottom*) with a schematic illustration demonstrating head angle direction for which the firing rate increases (*right top*). **(C)** Yaw angle and Yaw (L↑ R↓) neuron during horizontal tracking (*Pearson Correlation* (*PC*), *r*^2^ = 0.97, *p* < 0.0001). **(D)** Yaw angle and Yaw (L↓ R↑) neuron during horizontal tracking (*PC, r*^2^ = 0.89, *p* < 0.0001). **(E)** Roll angle and Roll (L↑ R↓) neuron during horizontal tracking (*PC, r*^2^ = 0.98, *p* < 0.0001). **(F)** Roll angle and Roll (L↓ R↑) neuron during horizontal tracking (*PC, r*^2^ = 0.98, *p* < 0.0001). **(G)** Pitch angle and Pitch (U↑ D↓) neuron during vertical tracking (*PC, r*^2^ = 0.98, *p* < 0.0001). **(H)** Pitch angle and Pitch (U↓ D↑) neuron during vertical tracking (*PC, r*^2^ = 0.99, *p* < 0.0001).

During reward tracking, mice showed a characteristic rotation of their head about three principal axes: yaw, roll, and pitch (Figure 1B, Videos S1, S2, and S3). During horizontal spout movement, mice significantly altered the roll and yaw angle of their head in a rhythmic fashion depending on the location of the spout (Videos S1 and S2), while the pitch angle largely stayed the same. On the other hand, during vertical spout movement, the pitch angle would characteristically fluctuate, but the roll and yaw angles remained relatively constant (Video S3). To understand the relationship between VTA GABAergic output and their pursuit behavior, we compared single unit activity with a number of recorded behavioral variables including the pitch, yaw, and roll angles (Figure S1). We discovered that the firing rates of many recorded neurons exhibited extremely high correlations with instantaneous head angles (Figures 1C-H). Strikingly, we were able to identify three distinct classes of VTA^Vgat+^ neurons corresponding to the yaw, pitch, and roll axes of rotation (Figure 1).

Because neurons in the VTA are difficult to classify based solely on spike waveform width and firing rate (Cohen et al., 2012; Margolis et al., 2006), we confirmed our classification using optotagging. We stimulated channelrhodopsin (ChR2) infected VTA^Vgat+^ neurons in *Vgat-ires-Cre* mice while recording their single-unit activity (*n* = 18, Figure 2). Results from our opto-tagging experiments revealed distinct waveforms compared to the remaining population. Using an unsupervised clustering algorithm based on the first 10 principal components of the average waveforms, we identified two separate clusters. One of these clusters contained all of the optically identified GABAergic neurons, and was subsequently used to identify the remaining GABAergic population (Figures 2F-H).

**Figure 2.**
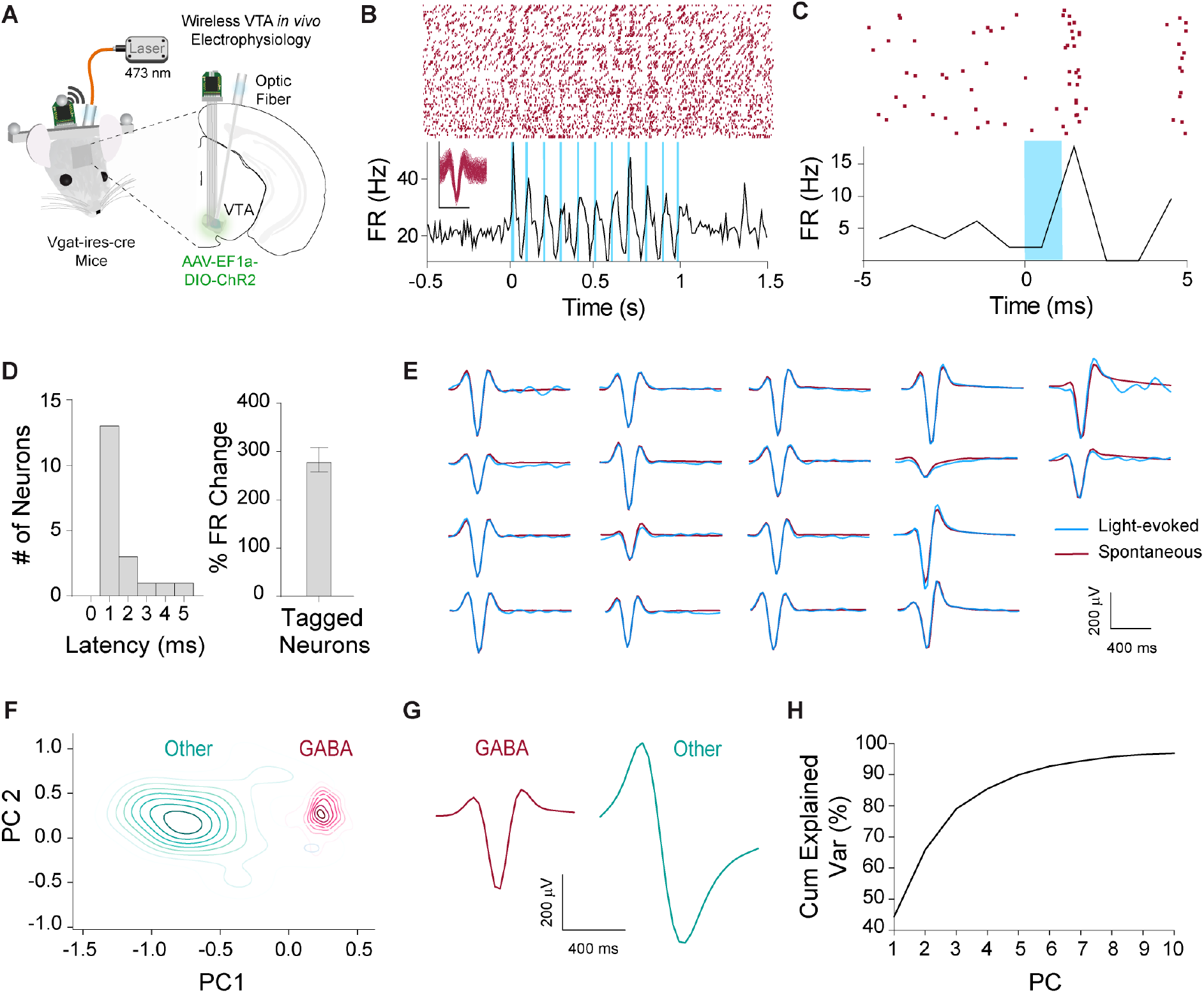
Optogenetic identification of VTA^Vgat+^ neurons. **(A)** Schematic illustration of optrode in the VTA for simultaneous optogenetic excitation and recording of *VGAT+* neurons in *Vgat-ires-Cre* mice. **(B)** Representative example of an optically tagged VTA^Vgat+^ neuron using 10 Hz stimulation. **(C)** Representative neuron showing a 1 ms latency between light stimulation and neural response. **(D)** *Left*: All tagged GABAergic neurons had a latency of < 5 ms (*n =* 18). *Right*: Optically tagged GABAergic neurons displayed a significant increase in neuronal firing activity compared to baseline (*p* < .0001). **(E)** Fidelity analysis demonstrating similarity of waveforms between spontaneous (maroon) and light-evoked (blue) waveforms. **(F)** Contour plot of the first two principal components out of 10 within an unsupervised clustering of neuronal waveforms. *Post hoc* analysis revealed all optically tagged VTA^Vgat+^ neurons fell within one cluster. The remaining neurons that fell within the same cluster were classified as putative GABAergic neurons. **(G)** Average waveforms for cells classified as GABAergic (*n* = 421) or other (*n* = 58). **(H)** Cumulative explained variance for each principal component. Ten principal components accounted for over 95% of the variance within the neuronal waveforms

For each neuron class, the firing rate varied monotonically with the head angle, and was selective for a single axis of rotation (Figures 1C-H). For example, a given neuron would increase its firing rate when the head was tilting to the left, and would decrease firing when tilting to the right (Figure 1E). For each axis of rotation, we found two opponent populations based on the direction of rotation (Figure 3; Figure S3). There are thus six distinct populations of VTA GABA neurons related to instantaneous head angle. Moreover, a neuron that represents one axis of rotation would show weaker correlation with rotation about the other axes (Figures 4A-C). These correlations were extremely robust, apparent even at a single trial level (Figure S2). For both roll and yaw angle neurons, we found more neurons whose firing rates increased in the ipsiversive direction relative to the recording hemisphere (Figures 4I-J). Because the yaw and roll angle behavior co-varied when the animals were following a horizontally moving target, we compared neurons classified as either VTA^Vgat+^ roll or VTA^Vgat+^ yaw. The average correlational value for neurons classified as VTA^Vgat+^ roll was significantly higher than its average correlational value with yaw angle (Figure 4B). This significant difference was also found for neurons classified as VTA^Vgat+^ yaw neurons with respect to roll angle (Figure 4C).

**Figure 3.**
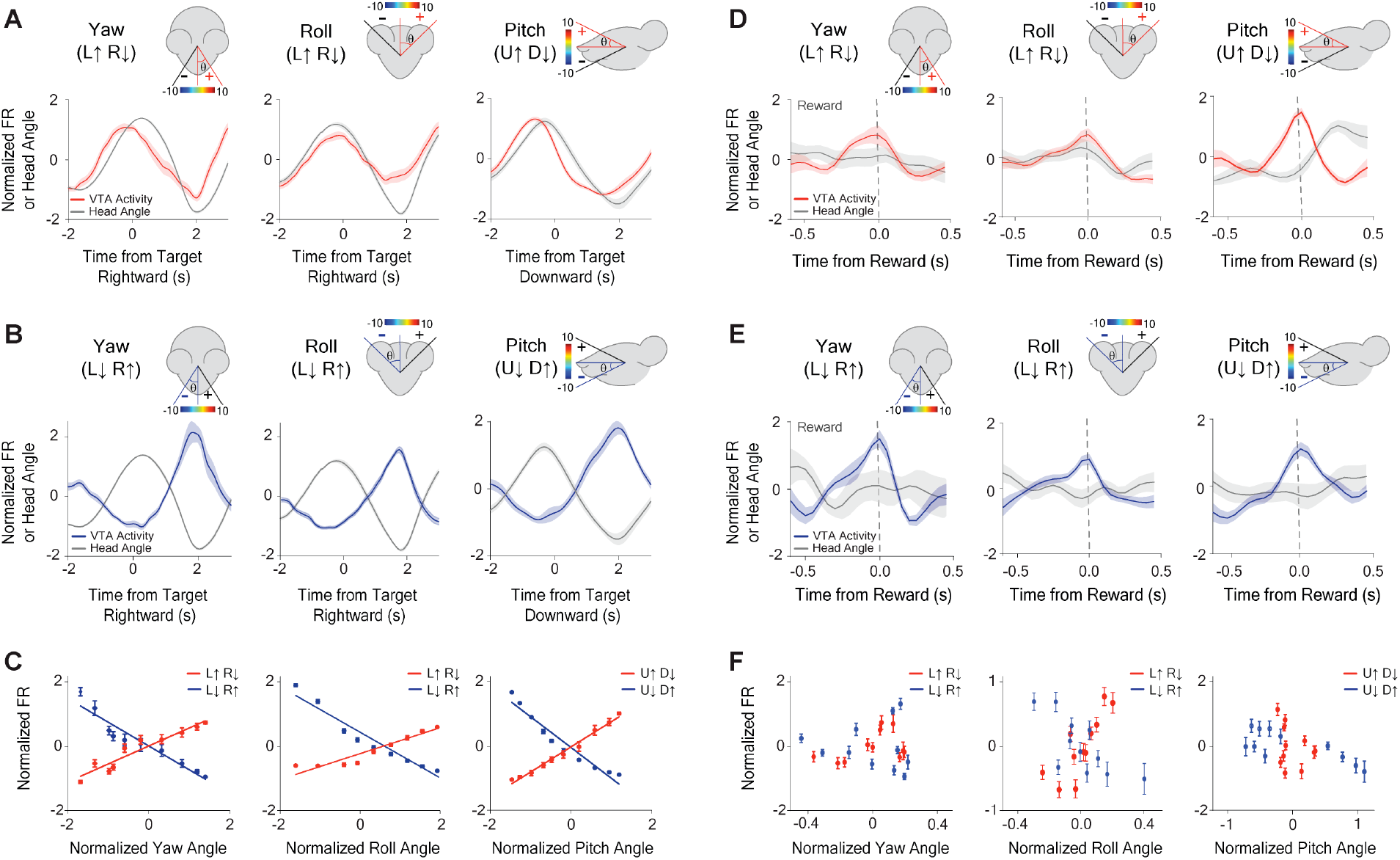
VTA GABAergic ensembles represent yaw, roll, and pitch, but these representations degrade at the time of reward. **(A-C)** Population activity precisely represents orthogonal angles of head rotation. **(A)** Neural population for Yaw (L↑ R↓) neurons (*left*, *n* = 16), Roll (L↑ R↓) neurons (*middle, n* = 28), and Pitch (U↑ D↓) neurons (*right*, *n* = 16) during reward tracking. Traces and error bars represent the Mean ± SEM. **(B)** Neural population for Yaw (L↓ R↑) neurons (l*eft*, *n* = 10), Roll (L↓ R↑) neurons (*middle*, *n* = 34), and Pitch (U↓ D↑) neurons (*right*, *n* = 13) during reward tracking. **(C)** Population average showing correlation between head angle and VTA neuronal firing rate while tracking reward: Yaw angle (*left*) (Yaw (L↑ R↓): *PC, r*^2^ = 0.91, *p* < .0001, *n* = 16; Yaw (L↓ R↑): *PC, r*^2^ = 0.92, *p* < 0.0001, *n* = 10). Roll Angle (*middle*) (Roll (L↑ R↓): *PC, r*^2^ = 0.92, *p* < 0.0001, *n* = 28; Roll (L↓ R↑): *PC, r*^2^ = 0.94, *p* < 0.0001, *n* = 34). Pitch Angle (*right*) (Pitch (U↑ D↓): *PC, r*^2^ = 0.99, *p* < 0.0001, *n* = 16; Pitch (U↓ D↑): *PC, r*^2^ = 0.94, *p* < 0.0001, *n* = 13). **(D-F)** Degradation of neural coding for head angle at the time of reward consumption. **(D)** Neural population for Yaw (L↑ R↓) neurons (*left*, *n* = 16), Roll (L↑ R↓) neurons (*middle, n* = 28), and Pitch (U↑ D↓) neurons (*right*, *n* = 16) at the time of reward. **(E)** Neural population for Yaw (L↓ R↑) neurons (l*eft*, *n* = 10), Roll (L↓ R↑) neurons (*middle*, *n* = 34), and Pitch (U↓ D↑) neurons (*right*, *n* = 13) at the time of reward. **(F)** Population average showing correlation between head angle and VTA neuronal firing rate during reward consumption: Yaw angle (*left*) (Yaw (L↑ R↓): *PC, r*^2^ = 0.18, *p* > 0.05 *n* = 16; Yaw (L↓ R↑): *PC, r*^2^ = 0.005, *p* > 0.05, *n* = 10). Roll Angle (*middle*) (Roll (L↑ R↓): *PC, r*^2^ = 0.65, *p* = 0.003, *n* = 28; Roll (L↓ R↑): *PC, r*^2^ = 0.53, *p* = 0.01, *n* = 34). Pitch Angle (*right*) (Pitch (U↑ D↓): *PC, r*^2^ = 0.09, *p* > 0.05, *n* = 16; Pitch (U↓ D↑): *PC, r*^2^ = 0.56, *p* = 0.01, *n* = 13). Traces and error bars represent the Mean ± SEM.

**Figure 4.**
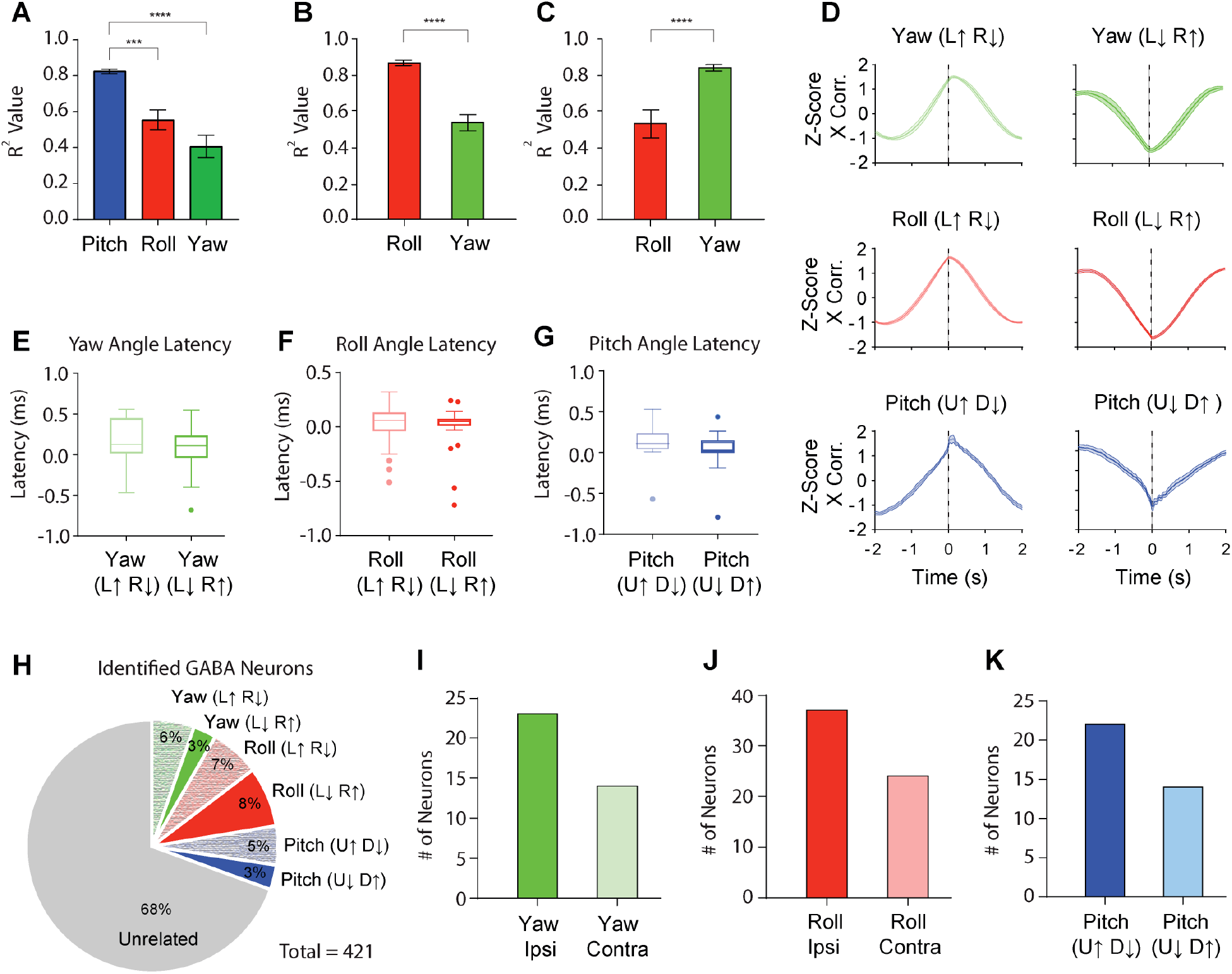
Summary of classified VTA GABAergic neurons. **(A)** Average *r*^2^ values for yaw, roll and pitch from a Pearson Correlation (*PC)* analysis for neurons classified as pitch. (Pitch: *r*^2^ = 0.83 ± 0.01; Roll: *r*^2^ = 0.56 ± 0.03; Yaw: *r*^2^ = 0.41 ± 0.05; Mean ± SEM). Neurons correlated with pitch angle are significantly different from roll and yaw angle (*F*_*(2,59)*_ = 19.52, *p* < 0.0001; *p* values were corrected with Dunnett’s multiple comparison test. *** *p* = 0.0003; **** *p* < 0.0001). **(B-C)**Because yaw and roll co-varied during horizontal tracking and had similar *r*^2^ values when compared to the pitch population, we directly compared *r*^2^ values between roll and yaw angle for neurons classified as yaw or roll angle neurons. **(B)** There is a significant difference between the *r*^2^ values of yaw and roll angle for neurons categorized as roll neurons (paired t-test, *t*_(61)_ = 10.57, *p* < 0.0001, Roll: *r*^2^ = 0.87 ± 0.01; Yaw: *r*^2^ = 0.54 ± 0.03; Mean ± SEM). **(C)** There is also a significant difference between the *r*^2^ values of yaw and roll angle for neurons categorized as yaw neurons (paired t-test, *t*_(25)_ = 6.26, *p* < 0.0001, Yaw: *r*^2^ = 0.83 ± 0.01; Roll: *r*^2^ = 0.54 ± 0.05; Mean ± SEM). **(D)** Normalized cross-correlations for yaw neurons (*top*), roll neurons (*middle*), and pitch neurons (*bottom*). Latency between neural activity and behavior were determined from maximum values of cross-correlations. **(E)** Latency between neural activity and yaw angle for both Yaw (L↑ R↓; 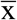 = 180 ± 0.05 ms) and Yaw (L↓ R↑; 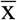 = 52 ± 0.09 ms; Mean ± SEM) neurons. **(F)** Latency between neural activity and roll angle for both Roll (L↑ R↓; 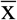 = 60 ± 0.03 ms) and Roll (L↓ R↑; (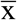 = 50 ± 0.03 ms; Mean ± SEM) neurons. **(G)** Latency between neural activity and pitch angle for both Pitch (U↑ D↓; 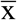 = 130 ± 0.04 ms) and Pitch (U↓ D↑; 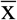 = 20 ± 0.07 ms; Mean ± SEM) neurons. **(H)** Breakdown of angle neuron percentages from VTA GABAergic neuron population. **(I)** Quantification of the number of yaw angle neurons that increase their firing rate in the ipsiversive direction (*n* = 23) or contraversive direction (*n* = 14) in relation to the recording hemisphere. **(J)** Quantification of the number of roll angle neurons that increase their firing rate in the ipsiversive direction (*n* = 37) or contraversive direction (*n* = 24) in relation to the recording hemisphere. **(K)** Quantification of the number of pitch angle neurons that increase their firing rate in the upward direction (*n* = 22) or downward direction (*n* = 14).

In order to examine whether we could predict head angle from all GABAergic neurons, not just those classified as head angle neurons, we used a machine learning algorithm (Support Vector Regression, SVR). With the SVR, we were able to decode actual head angle from all simultaneously recorded VTA GABAergic neurons (Figure S4).

Next, we examined the activity of these different neuron classes at the time of reward delivery. Interestingly, the correlations between the neural activity and head angle degraded at the time of reward. Independently of their angle representation or direction preference, they often show a transient increase at the time of reward delivery (Figures 3D-F; Figures S3G-I). The population response at the time of reward is similar to what was previously reported (Cohen et al., 2012).

Although our electrophysiological classification of GABAergic neurons was validated using optotagging experiments, we could not be sure that the remaining classified neurons were in fact GABAergic because our GABAergic population was uncharacteristically high for VTA GABAergic neurons. This could be due to the impedance characteristics of our electrodes, indistinguishable waveforms, or the higher firing rates of GABAergic neurons in relation to their neighboring dopamine neurons. Thus, to further confirm that these neurons are GABAergic, we injected the fluorescent calcium indicator GCaMP7f into the VTA of *Vgat-ires-Cre* mice and used a miniature 1-photon microscope for chronic *in vivo* imaging (Figure 5A) (Cai et al., 2016; Dana et al., 2018; Flusberg et al., 2008). This allowed us to simultaneously record calcium fluctuations (*n* = 158) and behavior during reward-tracking. Consistent with our electrophysiological data, we found GABAergic neurons that were correlated with yaw, roll, and pitch (Figure 5; Figure S5). In addition, we found more yaw and roll neurons that increased their firing rates in the ipsiversive direction relative to the hemisphere being imaged (Figures S6C-D).

**Figure 5.**
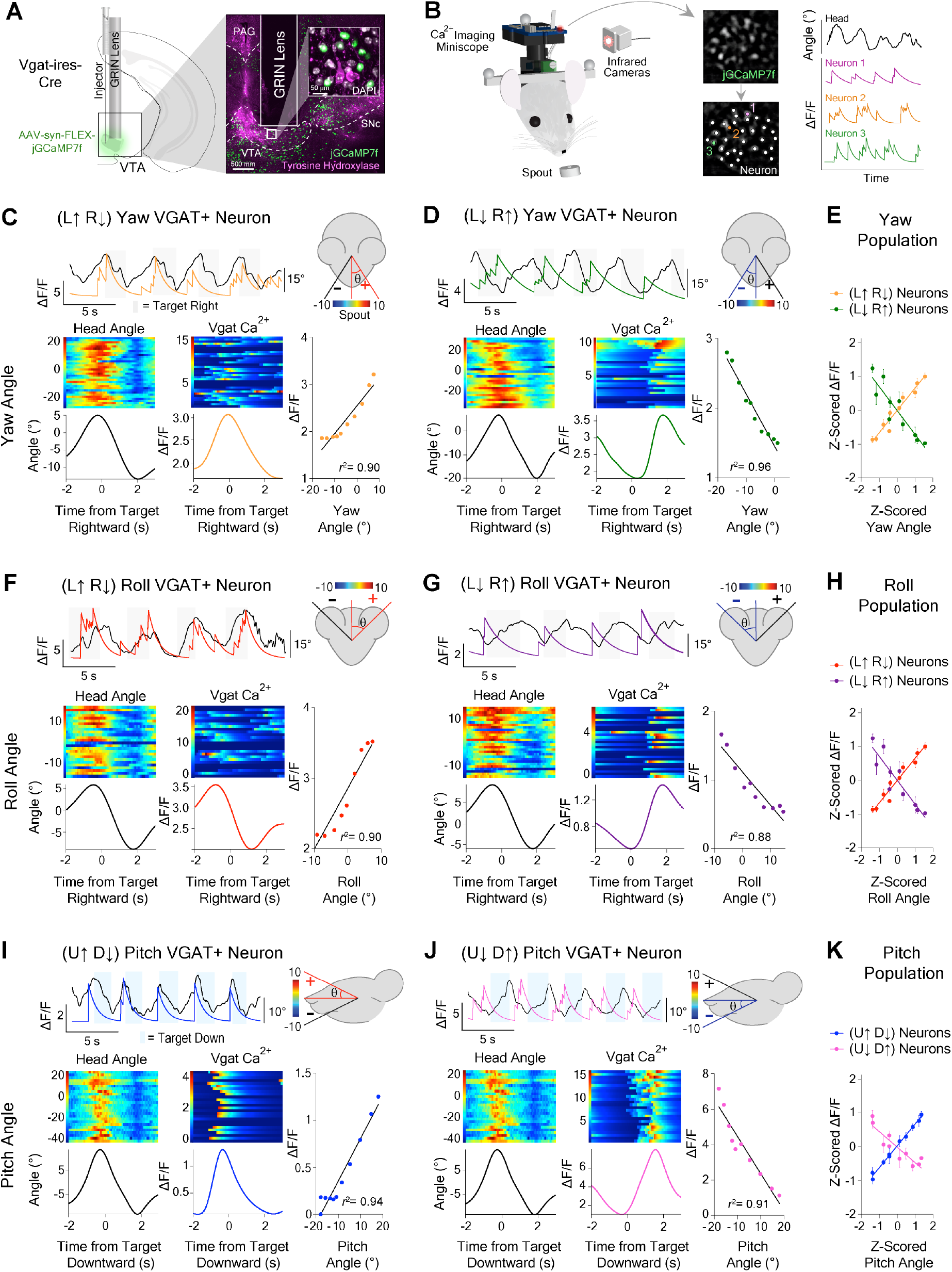
Confirmation of orthogonal head angle representations using *in vivo* calcium imaging of VTA GABAergic neurons. **(A)** GRIN lens implantation over AAV-hSyn-GCaMP7f infected GABAergic cells in the VTA of *Vgat-ires-Cre* mice for chronic *in vivo* recordings of calcium fluctuations. Representative coronal section through the midbrain shows calcium indicator (jGCaMP7f) expression in the vicinity of tyrosine hydroxylase neurons in the VTA. **(B)** Schematic illustration of 1-photon miniscope attached to mouse head for calcium imaging during reward tracking behavior (*left*). Extracted changes in fluorescent intensity of VTA^Vgat+^ neurons concomitant with rotational head kinematics (*right*). **(C-J)**VTA^Vgat+^ neurons represent head angles about three orthogonal axes of rotation. Raw calcium transients and angle behavior over 20s (*top*). Black traces represents head angle, colored traces represent calcium transients. Peri-event heat maps of head angle (*left*) and calcium activity (*middle*). Correlation between neural activity and angle (*right*). **(C)** Yaw (L↑ R↓) neuron (*PC*, *r*^2^ = 0.90, *p* < 0.0001). **(D)** Yaw (L↓ R↑) neuron (*PC*, *r*^2^ = 0.96, *p* < 0.0001). **(E)** Correlation for Yaw angle population (Yaw (L↑ R↓): *PC, r*^2^ = 0.95, *p* < 0.0001, *n* = 18; Yaw (L↓ R↑): *PC, r*^2^ = 0.83, *p* = 0.0002, *n* = 14). **(F)** Roll (L↑ R↓) neuron (*PC*, *r*^2^ = 0.90, *p* < 0.0001) **(G)** Roll (L↓ R↑) neuron (*PC*, *r*^2^ = 0.88, *p* < 0.0001). **(H)** Correlation analyses for Roll angle population (Roll (L↑ R↓): *PC, r*^2^ = 0.94, *p* < 0.0001, *n* = 16; Roll (L↓ R↑): *PC, r*^2^ = 0.65, *p* = 0.005, *n* = 14). **(I)** Pitch (U↑ D↓) neuron (*PC*, *r*^2^ = 0.94, *p* < .0001). **(J)** Pitch (U↓ D↑) neuron (*LR*, *r*^2^ = 0.91, *p* < 0.0001). **(K)** Correlation for Pitch angle population (Pitch (U↑ D↓): *PC, r*^2^ = 0.99, *p* < 0.0001, *n* = 15; Pitch (U↓ D↑): *PC, r*^2^ = 0.82, *p* = 0.0003, *n* = 12).

VTA neural activity usually preceded head angle change (Figures 4D-G, Yaw: 117 ± 54 ms; Roll: 55 ± 23 ms; Pitch: 76 ± 29 ms). This suggests that VTA^Vgat+^ neurons send top down commands to generate movements with the specified rotational kinematics. To test this hypothesis, we used optogenetics to manipulate VTA^Vgat+^ neurons, using channelrhodopsin (ChR2) for excitation or soma-targeted *Guillardia theta* anion-conducting ChR2 (stGtACR2) for inhibition (Figure 6A; Vgat::ChR2^VTA^, *n* = 6; Vgat::stGtACR2^VTA^, <u>*n*</u> = 6; control, Vgat::eYFP^VTA^, *n* = 7) (Boyden et al., 2005; Mahn et al., 2018). We found that both optogenetic excitation and inhibition of VTA neurons produce opposite rotations of the head in all three axes (Figures. 6C-D, Figure S7). During excitation, the head is lowered (pitch angle decreased relative to the longitudinal axis of the body), and the roll and yaw angles deviate towards the stimulated hemisphere (Figures 6C-D; Video S4). There is therefore a predominantly ipsiversive effect on head angle. During inhibition, the head is raised (increased pitch angle), and the roll and yaw angles deviated away from the stimulation hemisphere, showing a contraversive effect (Figures 6C-D, Video S5). These results agree with our electrophysiology and imaging results. Strikingly, parametric manipulations of pulse width quantitatively determined head angle (Figure 6C). The latency between stimulation and behavioral changes was approximately the same for each axis of rotation (Figures S7D-F, Yaw: 37 ± 5 ms; Roll: 37 ± 4 ms; Pitch: 39 ms ± 6 ms).

**Figure 6.**
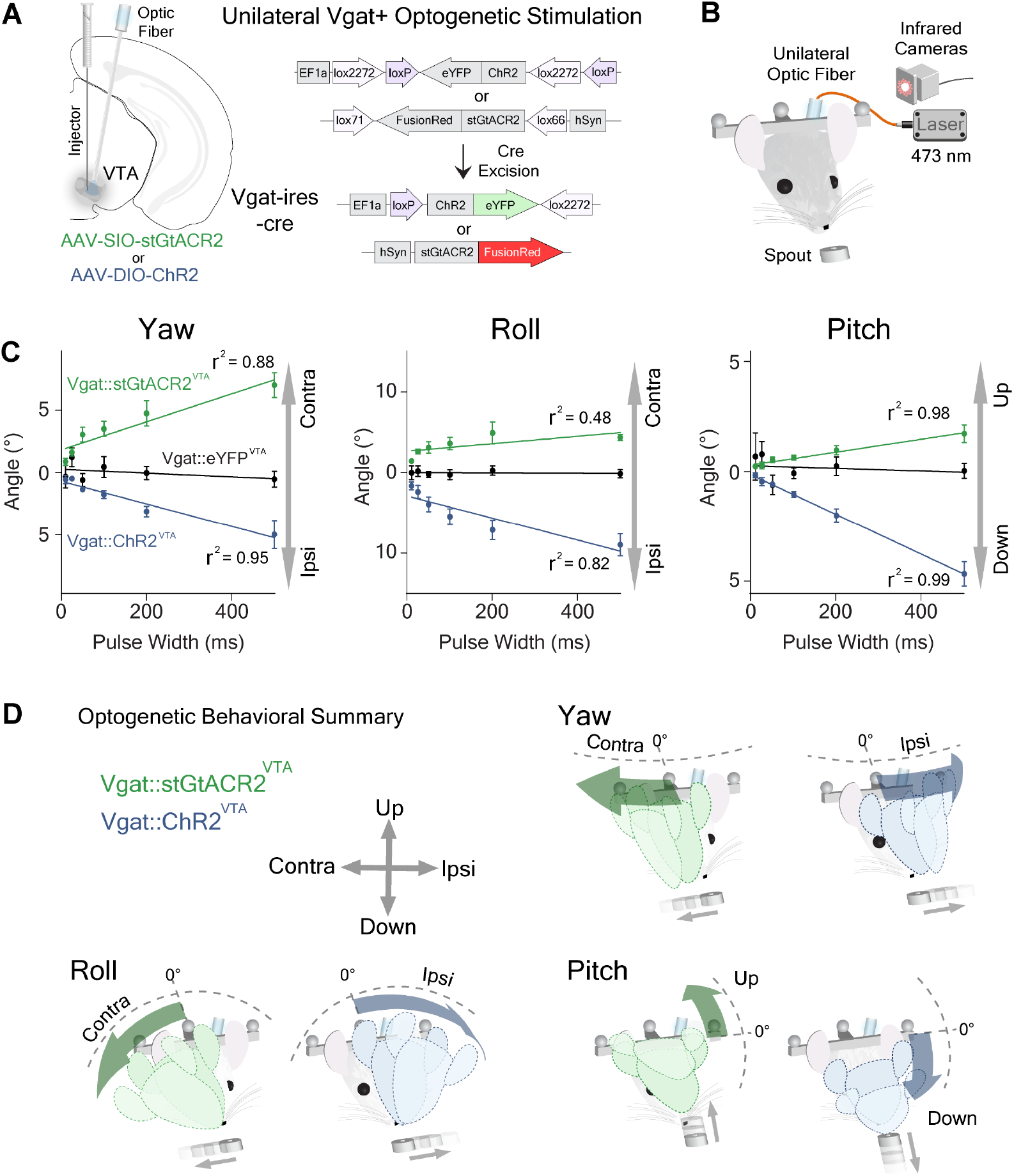
Optogenetic excitation and inhibition produce opposite deviations in head angles. **(A)** Unilateral stimulation of VTA^*Vgat+*^ neurons. AAV-DIO-ChR2 or AAV-SIO-StGtACR2 was injected into the VTA of *Vgat-ires-Cre* mice. **(B)** Schematic of unilateral optogenetic stimulation during reward tracking. **(C)** Optogenetic relationship between pulse width and deviation of pitch (*left*), roll (*middle*) and yaw (*right*) angles (*two-way RM* ANOVA, Yaw Angle, Main effect of Group, *F*_*(2,17)*_ = 91.93, *p* < 0.0001; Roll Angle, Main effect of Group, *F*_*(2,167)*_ = 72.04, *p* < 0.0001; Pitch Angle, Main effect of Group, *F*_*(2,17)*_ = 35.63, *p* < 0.0001, ChR2: *n* = 6; StGtaCR2: *n* = 6; control: *n* = 7). **(D)** Schematic illustration of the effects of stimulation on yaw, roll, and pitch. Optogenetic excitation produces ipsiversive and downward deviations of head angles, while inhibition produces contraversive and upward deviations of head angles.

Furthermore, selective optogenetic excitation of VTA^Vgat+^ neurons reduced reward consumption, as previously reported (van Zessen et al., 2012). This appeared to be caused by an increase in the distance between their head and the spout during stimulation (Figure 7H). Using our stimulation parameters, there did not appear to be any aversive reactions. We confirmed this in a real-time place preference assay, in which entering one side of the arena produces sustained optogenetic stimulation of the VTA^Vgat+^ neurons (Figure 7).

**Figure 7.**
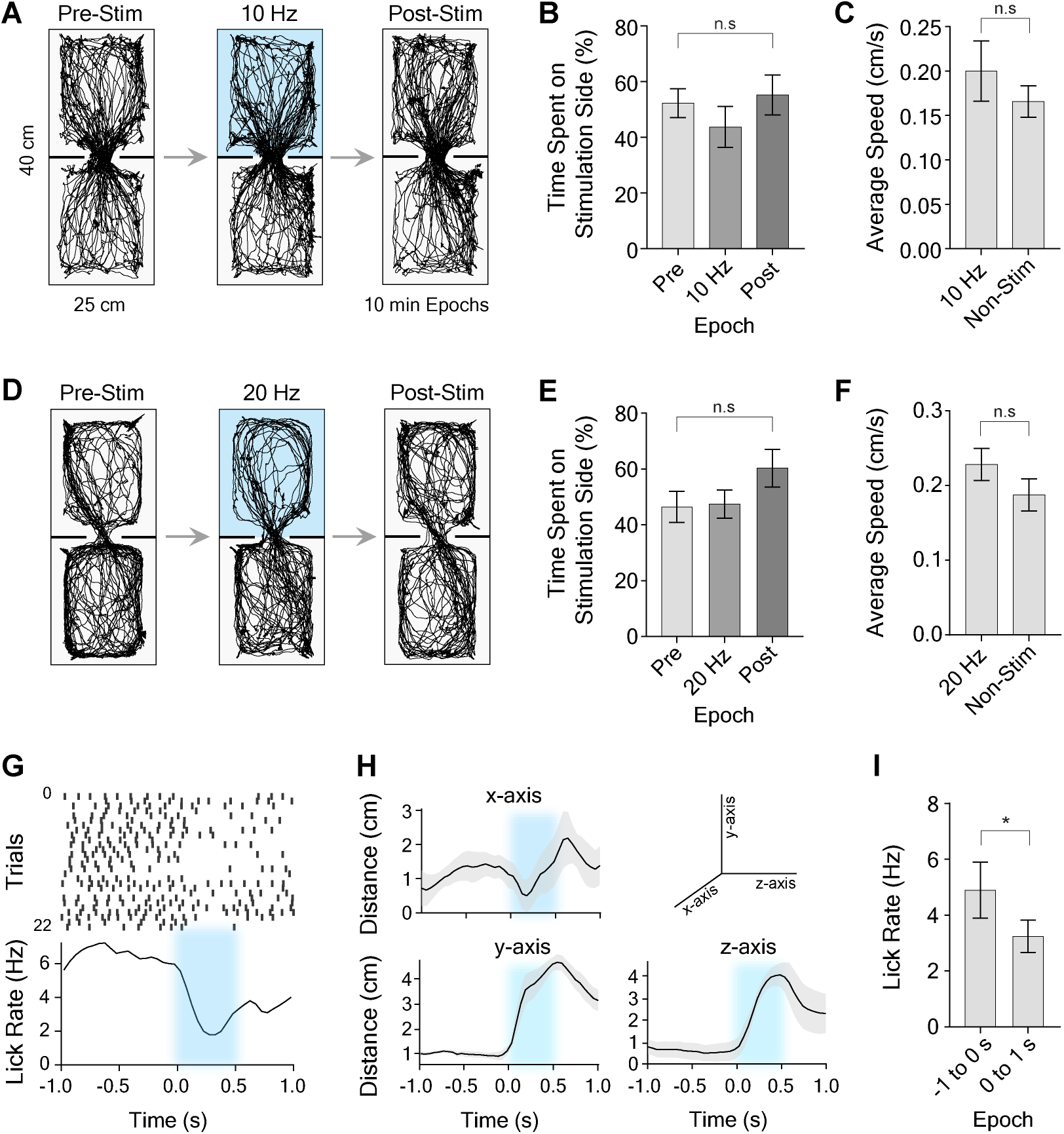
VTA GABAergic stimulation has no effect in real-time conditioned place preference, but reduces reward consumption in reward-tracking task. **(A)** Raw traces during real-time conditioned place preference assay using 10 Hz stimulation (10 ms pulse width). Mice were placed into a chamber for a total of 30 minutes across three epochs (pre-stimulation, stimulation, post-stimulation epochs; 10 min per epoch). During the stimulation epoch, 10 Hz unilateral stimulation occurred when the animal was in the top half of the chamber. **(B)** There was no significant difference between the time spent on the 10 Hz stimulation side versus the pre- and post-stimulation epochs (RM one-way ANOVA, *F*_*(2,21)*_ = 3.99, *p* = 0.01, *n* = 8. *Post-hoc* analysis that corrected for multiple comparisons, however, revealed no significant differences between groups, *p* > 0.05). **(C)** Average speed during 10Hz stimulation epoch on the stimulation half of the chamber was not significantly different than the non-stimulation half (paired t-test, *t*_(7)_ = 0.89, *p* > 0.05, *n* = 8). **(D)** Raw traces during real time-CPP during 20 Hz stimulation. **(E)** There was no significant difference between the time spent on the 20 Hz stimulation half of the chamber (10 ms pulses versus the pre- and post-stimulation epochs; RM one-way ANOVA, *F*_*(2,21)*_ = 3.48, *p* = 0.02, *n* = 8. *Post-hoc* analysis that corrected for multiple comparisons revealed no significant differences between groups, *p* > 0.05). **(F)** Average speed during 20 Hz stimulation epoch on the stimulation side of the chamber was not significantly different than the non-stimulation side (paired t-test, *t*_(7)_ = 0.09, *p* > 0.05, *n* = 8). **(G-I)**Licking-related activity during reward tracking task. **(G)** Representative licking raster during 500 ms of optogenetic excitation. Top panel is a peri-event raster plot, where each dash represents a lick, and each row represents one trial. Bottom panel is average lick rate plot. Excitation produced a reduction in licking during and immediately after licking. **(H)** Population graphs showing increased distance to the target due to stimulation for all three axes (*n* = 6). **(I)** Average lick rate was significantly reduced during and immediately after stimulation compared to before stimulation (paired t-test, *t*_(5)_ = 3.20, *p* =0.02, *n* = 6).

## Discussion

Collectively, our results show for the first time that VTA^Vgat+^ neurons determine rotational kinematics of the head, and shed light on the computational role of these neurons. The major advantage of our approach is the lack of head restraint, so that the relationship between neural activity and free behavior is revealed for the first time. We found three distinct classes: the instantaneous firing rate of each represents a separate angle (pitch, roll and yaw). Furthermore, each class can be further divided into populations that increase firing in a given head tilt direction. It appears that each population is primarily responsible for tilt in a single direction. In order to achieve bi-directional control within the system, opponent populations are needed to increase or decrease a particular angle about a particular axis of rotation. Our findings suggest that VTA GABAergic neurons not only represent head angles, but also generate desired angles during continuous behavior. This is possible if their output, acting as reference signals, quantitatively dictate the sensory states to be reached by lower level systems (Yin, 2017).

While our results demonstrate the role of VTA GABAergic neurons in control of head orientation, it does not follow that this is their sole function. Much like their neighboring DA neurons, GABA neurons receive a diverse set of inputs and have wide-ranging connections to many different brain regions, suggesting that they may have multiple functional roles in learning and behavior (Beier et al., 2019; Lammel et al., 2012; Su et al., 2019; Tan et al., 2012; Yu et al., 2019). Interestingly, their representation of head angle is transiently degraded at the time of reward (Figures 3D-F), and the GABAergic neurons usually increase firing at the time of reward, regardless of their head angle coding.

Given the dominant view that the VTA is crucial for motivated behaviors, one obvious question is how the present findings could be related to the extensive literature on appetitive and aversive behaviors. A few studies have examined the role of the VTA GABAergic neurons in behavior using cell-type specific manipulations. However, because they either relied on head-fixed preparations, or used behavioral measures such as place preference, average velocity, or total distance, they were not able to show the coordination of head rotation kinematics we demonstrate here (Tan et al., 2012; van Zessen et al., 2012).

Previous studies found that stimulation of VTA GABAergic neurons interrupted reward consumption (van Zessen et al., 2012). This is not surprising given our finding that optogenetic manipulation of these neurons predictably resulted in head deviation. Even in head-fixed mice, it is possible that such deviations are sufficient to interrupt reward consumption. We also found that consumption was reduced during optogenetic stimulation, primarily due to stimulation-induced head rotation, which increases distance from the reward spout (Figure 7). Previous work also suggests that, by inhibiting DA neurons, VTA GABAergic neurons could perform a subtraction needed for calculation of prediction errors(Eshel et al., 2015). While our results do not rule out any computational role in prediction errors, they show for the first time that these neurons directly command the actual behavior in a spatially precise manner (Figure 6).

Our results also suggest that a separate process mediating the consummatory phase of behavior may take over and command VTA GABAergic neurons at the time of reward. This may explain why reward-related activity is often seen in these neurons, and why the correlations with head angle seem to disappear at the time of reward receipt (Figures 3D-F; Figures S3G-I). The control processes that mediate the transition from appetitive to consummatory phases of motivated behaviors must be precisely coordinated and overlap in time to achieve smooth control. Additional studies will be needed to elucidate the circuits underlying tracking and consummatory aspects of the behavior.

While head rotation control is absolutely essential for most motivated behaviors, it has largely been neglected in previous studies. Brain systems for precise orienting and steering are necessary for nearly all aversive and appetitive behaviors, whether seeking reward or avoiding harm. For example, in consummatory behavior like eating, the mouse must make continuous and subtle adjustments of the head. There is therefore no need to dissociate head orientation control from reward and aversion. Excessive vestibular disturbance, for example, can be highly aversive and can result in disorientation and nausea. Indeed, during our optogenetic experiments, the mice also sometimes showed urination and defecation in addition to the head angle changes. While our results did not show aversive effects (Figure 7), this could explain why others have found that optogenetic excitation of VTA GABAergic neurons is aversive (Tan et al., 2012). It is also important to note that our stimulation parameters are more physiological (10-20 Hz vs. constant stimulation) than those used in previous work, which could also explain the lack of a clear aversive effect.

A complex and multi-level circuitry is required to implement fine control of the head in space, given the multiple degrees of freedom involved. Our results suggest that the brain solves this control problem using a similar strategy as in airplanes or ships, using independent control of three orthogonal axes of rotation. Distinct neuronal populations in the VTA are responsible for independent control of rotation along these axes. The observation that activity in these cells generally leads the kinematic variable suggests that VTA GABAergic neurons do not simply provide perceptual representations of head angle. The signals are not predictions of future perceptual signals, but commands that dictate the perceptions to be reached, as demonstrated by our optogenetic stimulation results. The amount of stimulation quantitatively determines the head angle achieved. In normal movements, all the populations are engaged, though to different extents depending on the degree of rotation along each axis.

Interestingly, the VTA is directly connected to key nuclei in the head direction circuit (Angelaki and Yakusheva, 2009; Moser et al., 2008; Taube, 2007). Angular head velocity cells have been reported in the dorsal tegmental nucleus, superior colliculus, and the lateral habenula, which are all connected to the VTA (Bassett and Taube, 2001; Faget et al., 2016; Sharp et al., 2006; Taube, 2007; Taylor et al., 2014; Wilson et al., 2018). Thus, VTA GABAergic neurons could be integrating signals representing angular head velocity into accurate representations of head angle. The interactions between the VTA and the other components of the head steering circuit remain to be investigated.

## References and Notes

## Acknowledgments

We would like to thank Dr. Fengxia Allen, Dr. Guozhong Yu, Dr. Joseph Barter, Dr. Jinyong Zhang, Lee Christensen, and Murray Wickwire for their technical assistance. We would like to thank Dr. Joseph Barter for comments and discussions. We also thank Dr. Nicole Calakos and Dr. Brandon Turner for providing the gCaMP7f virus.

## Author contributions

R.N.H. and H.H.Y. conceptualized and designed studies. R.N.H performed surgeries, behavioral experiments, electrophysiological recordings and calcium imaging. G.D.R.W. performed immunohistochemistry, confocal imaging, and helped design figures. N.K developed the tracking task. N.K, K.I.B., E.A.P., and R.N.H. wrote data collection and analysis code. K.I.B. captured and edited videos. R.N.H. and H.H.Y. drafted the manuscript. K.I.B., G.D.R.W., E.A.P., R.N.H. and H.H.Y edited and revised the manuscript. All authors have read and approved the manuscript.

## Data and materials availability

All data are available in the main text or the supplementary materials. Raw data available upon request from corresponding author

## Supplementary Materials

Materials and Methods

Supplementary Figures 1-8

Supplemental Videos 1-5

