## Supplemental for "Precise Coordination of 3-dimensional Rotational Kinematics by Ventral Tegmental Area GABAergic Neurons"

**This PDF file includes:**

STAR Methods
Supplementary Figures S1 to S8
Captions for Movies S1 to S5

**Other Supplementary Materials for this manuscript include the following:**

Movies S1 to S5

**STAR Methods**

All experimental procedures were conducted in accordance with standard ethical guidelines and were approved by the Duke University Institutional Animal Care and Use Committee.

**Contact for Reagents and Resource Sharing**

**Key Resources Table**

| ***REAGENT or RESOURCE*** | ***SOURCE*** | ***IDENTIFIER*** |
|---|---|---|
| ***Antibodies*** | | |
| Goat polyclonal anti-Rabbit Alexa Fluor 488 | abcam | Cat# ab150077; RRID: AB_2630356 |
| Goat polyclonal anti-Rabbit Alexa Fluor 594 | abcam | Cat# ab150080; RRID: AB_2650602 |
| Rabbit polyclonal anti-Dopamine Transporter | abcam | Cat# ab111468; RRID: AB_11155293 |
| Rabbit polyclonal anti-Tyrosine Hydroxylase | Millipore | Cat# 657012; RRID: AB_566341 |
| Rabbit polyclonal anti-Vesicular GABA Transporter | Millipore | Cat# AB5062P; RRID: AB_2301998 |
| ***Bacterial and Virus Strains*** | | |
| rAAV5-EF1α-DIO-eYFP | Duke Vector Core | N/A |
| rAAV5-EF1α-DIO-hChR2(H134R)-eYFP | Duke Vector Core | N/A |
| pGP-AAV1-syn-jGCaMP7f-WPRE | addgene | Cat# 10448-AAV1 |
| AAV1-hSyn-SIO-stGtACR2-FusionRed | addgene | Cat# 105677-AAV1 |

| | | |
|---|---|---|
| ***Experimental Models: Organisms/Strains*** | | |
| Mouse: Vgat-ires-Cre: $1^{tm2(cre)Lowl}$/J | Jackson Laboratory | Mouse Strain: 016962 |
| Mouse: C57BL/6J | Jackson Laboratory | Mouse Strain: 000664 |
| ***Oligonucleotides*** | | |
| Primers for wild type mouse Vgat-ires-Cre<br><br>(F: GGTCGATGCAACGAGTGATGAGG)<br><br>(R: GCCAGATTACGTATATCCTGGCAG) | Integrated DNA Technologies | N/A |
| ***Software and Algorithms*** | | |
| MATLAB 2016b | MathWorks | https://www.mathworks.com/products/new_products/release2015b.html |
| Python 2.7 | Anaconda | https://www.anaconda.com/download/?lang=en-us |
| Bonsai | Open Ephys | http://www.open-ephys.org/bonsai/ |
| Offline Sorter 3.0 | Plexon | https://plexon.com/products/offline-sorter/ |
| NeuroExplorer 4.0 | Nex Technologies | http://www.neuroexplorer.com/downloadspage/ |
| Cortex 5.0 | MotionAnalysis | http://ftp.motionanalysis.com/html/industrial/cortex.html |
| GraphPad Prism 8 | GraphPad | https://www.graphpad.com/scientific-software/prism/ |

| *Other* | | |
|---|---|---|
| LED Driver | Thorlabs | LEDD1B |
| DAPI Fluoromount-G | Southern Biotech | Cat# 0100-20 |

**Animals:** All experimental procedures were approved by the Animal Care and Use Committee at Duke University. Both male and female C57BL/6J and *Vgat-ires-Cre* mice (3-8 months) acquired from Jackson Laboratory were used. Mice were maintained on a 12:12 light cycle and tested during the light phase. Mice used for *in vivo* electrophysiology and calcium imaging experiments were singly housed. All other mice were group housed. During behavioral experiments, mice were placed on water restriction. After training sessions, mice had free access to water for approximately 2 hours and were maintained at approximately 85-90% of their initial weights.

**Viral Constructs:** rAAV5.EF1α.DIO.hChR2(H134R).eYFP and rAAV5.EF1α.DIO.eYFP were obtained from the Duke University Vector Core. pAAV-hSyn1-SIO-stGtACR2-FusionRed (Mahn et al., 2018) and pGP-AAV-syn-jGCaMP7f-WPRE (Dana et al., 2018) were obtained from Addgene. pGP-AAV-syn-jGCaMP7f-WPRE was from Douglas Kim (Addgene viral prep # 104488-AAV1; http://n2t.net/addgene:104488 ; RRID:Addgene_104488). pAAV_hSyn1-SIO-stGtACR2-FusionRed was from Ofer Yizhar (Addgene viral prep # 105677-AAV1; http://n2t.net/addgene:105677; RRID:Addgene_105677).

**Surgery:** Mice were anesthetized with 2.0 to 3.0% isoflurane mixed with 0.60 L/min of oxygen for surgical procedures and placed into a stereotactic frame (David Kopf Instruments, Tujunga,

CA). Meloxicam (2 mg/kg) and topical bupivacaine (0.20 mL) were administered prior to incision. C57BL/6J mice (*n* = 26; 14 males, 12 females) were used for electrophysiology experiments. 16 channel electrode arrays (4x4 fixed or drivable) were lowered into the VTA (AP: 3.2 – 3.4 relative to bregma, ML: 0.4 – 0.6 relative to bregma, DV: 4.0 – 4.4 relative to brain surface) at a rate of 300 μm/min and grounded to a cranial screw. For optrode experiments, 200 nL of DIO-ChR2 was unilaterally injected into the VTA (AP: 3.2 – 3.4 relative to bregma, ML: 0.4 – 0.6 relative to bregma, DV: 4.0 – 4.4 relative to brain surface) of *Vgat-ires-Cre* mice using a microinjector (Nanoject 3000, Drummond Scientific) at a rate of 1 nL/s. The pipette was left to sit for 10 minutes at the injection site to allow absorption of the virus and prevent leakage. Optrodes were 4x4 microwire arrays (Neurophysiology Instruments) with a custom-made optic fiber positioned at an angle on the long side of the connector to maximize the number of neurons stimulated under the electrode array (Sparta et al., 2012). For optogenetic stimulation experiments, 200-300 nL of either AAV-DIO-ChR2 for excitation or AAV-SIO-stGtACR2 for inhibition was bilaterally injected into the VTA of *Vgat-ires-Cre* mice (*n* = 23; 11 males, 12 females) using the coordinates previously described. Custom-made optic fibers (5 - 6 mm length below ferrule, >80% transmittance, 105 μm core diameter) were then implanted directly above the VTA at an angle (AP: 3.2 – 3.4 with respect to bregma, ML: 1.6 with respect to bregma, DV: 3.8 from the brain surface; 15°). Fibers and electrodes were secured in place with dental acrylic adhered to skull screws. For calcium imaging experiments, *Vgat-ires-Cre* mice (*n* = 7; 3 males, 4 females) were unilaterally injected with 400 nL of AAV1-syn-jGCaMP7f-WPRE into the VTA at the coordinates and injection parameters previously described. A graded-index (GRIN) lens (Inscopix, 0.6mm x 7.1mm) was then slowly implanted above the VTA (AP: -3.2 – 3.5 with respect to bregma, ML: 0.4 – 0.8 with respect to bregma, DV: 4.0 from the brain surface) over the course of an hour. The lens was secured to implanted cranial screws with dental cement and covered with Kwik-Sil silicone elastomer (World Precision Instruments) to protect the surface of the lens. All mice were fitted with either a 3D printed headbar or custom-made steel headbar that could hold infrared tracking markers. Both heterozygous and homozygous *Vgat-ires-Cre* mice were used. Mice were allowed to recover for two weeks after surgery before experimentation.

**Wireless *in Vivo* Electrophysiology:** For electrophysiological recordings, either 4x4 16-channel fixed electrode ($n$ = 23) or drivable 16-channel arrays ($n$ = 3) (Innovative Neurophysiology, Inc.) were used. Driveable electrodes were moved 50-100 µm each session. If neurons in the same channel displayed a significantly similar waveform and ISI distribution after electrodes movement, they were excluded from analysis. Fixed arrays were constructed of tungsten wires (35 µm diameter, 150 µm spacing, 4 - 5 mm length). Drivable electrodes were single-drive movable micro-bundles (1 x 16) with 23 µm diameter electrodes driven through a guide cannula. VTA neural recordings were performed as previously described (Fan et al., 2011). Briefly, a miniaturized wireless head stage (Triangle Biosystems) or a tethered headstage interfaced with a Cerebrus data acquisition system (Blackrock Microsystems) was used to record activity during freely moving behavior. Electrophysiology Data were filtered with both analog and digital bandpass filters (analog highpass 1$^{st}$ order Butterworth filter at 0.3 Hz, analog lowpass 3$^{rd}$order Butterworth filter at 7.5 kHz, digital highpass 4$^{th}$ order Butterworth filter at 250 Hz). Filtered data was then sorted offline and analyzed using OfflineSorter (Plexon)
and Neuroexplorer (Nexus). The signal must have had a 3:1 signal-to-noise ratio, a refractory

period of at least 800 μs and consistent waveforms throughout the session to be considered as neural data for analysis.

**Optogenetic Stimulation and 3D Motion Tracking:** *Vgat-ires-cre* mice were used for optogenetic stimulation experiments. Unilateral stimulation data were taken from the same bilaterally implanted animal on separate experimental days. The output from the optic fiber tip was measured (PM120VA, ThorLabs) before each experimental session to obtain a power between 10-12 mW (i.e. ~10mW power delivered to the stimulation site with a transmittance of ~85%). A MATLAB program interfaced to a National Instruments DAQ triggered square pulse with varying durations (5 ms – 500 ms). Head movements were captured at 100 Hz in a Cartesian plane with eight Raptor-H digital infrared cameras (Motion Analysis, CA). To calculate the deviation of the head angle for pitch, roll, and yaw due to optogenetic stimulation, the angle of the head was first obtained at 100 ms prior to stimulation. This angle was then directly compared to the asymptote of the deviation immediately after simulation, or 200 ms after stimulation ended. For latency between stimulation and head angle changes, a 500 ms baseline was averaged together, and the latency was taken when a significant change within a 95% confidence window was reached.

**Behavioral Tasks and Analysis**:

*Reward Tracking Task*: Mice were placed on a custom-built platform (40 cm tall) and the platform was placed several cm away from a spout that was controlled by a stepper motor (Bipolar, 56.3 x 56.3 mm, DC 1.4A, 2.9Ω, 1.8 degree / step, Oriental motor, USA). To track movement, two infrared reflective markers (B & L Engineering) were attached to each side of a custom-made head bar attached to the dental cement skullcap of mice. A third marker was attached to either the top of the wireless electrode head stage, the top of the miniscope, or on a third bar attached to the skullcap. The reward position was tracked by placing a reflective marker approximately 20 mm away from the reward spout. Eight Raptor-H Digital infrared cameras were used to capture the movement of reflective markers during tracking. The data was then processed in Cortex (Motion Analysis, CA) and converted into Cartesian coordinates. Both reward delivery and the movement of the reward spout were controlled using custom MATLAB scripts. The program would deliver a 10% solution of condensed milk (12 µl every 500 ms) if mice tracked the moving spout (5-50 mm/sec) within a narrow window of Cartesian coordinates (X axis: 30 mm, Y axis: 20 mm, Z axis: 30 mm from the reward spout marker). To calculate roll and yaw angles, dimensions were reduced to either the y- or z-axis. Distance between markers (one on each side of the head, and one on top of the head) allowed calculation of the adjacent length of a triangle. Once all vector coordinates were obtained for one dimension, change in marker position was computed for each time step, providing the opposite length of the triangle. Because the distance between markers is the adjacent length, the arc tangent was then computed and converted into degrees for instantaneous angles using a custom MATLAB script. For pitch angle, the two side head bar markers and the third top marker created a triangle in 3D Cartesian space. Given the midpoint between two markers, the MATLAB function that converts Cartesian coordinates to spherical coordinates (car2sph) was used to compute the instantaneous pitch angle.

*Real Time Conditioned Place Preference*: Mice were placed on one side of a rectangular open field arena (45 cm L, 25 cm W, 35 cm H) that was partitioned in the middle by two 8 cm wide walls with a 9 cm opening. Videos were recorded with a camera that was placed 0.7 meters above the chamber and connected to a computer. Two-dimensional coordinates based on the center of mass of each mouse were collected at 50 f/s. An open source software program (Bonsai) using a custom script tracked movements for 30 minutes (Lopes et al., 2015). A session consisted of 3 separate, 10 min epochs: a pre-stimulation epoch where no stimulation occurred, a stimulation epoch where unilateral stimulation occurred only if the mice were on one half of the arena, and a post-stimulation epoch of no stimulation. Stimulation was produced using a laser attached to two Arduinos: one served as a data acquisition board to receive commands from the computer, and the other controlled the frequency output of the laser using TTL pulses. Data were analyzed using a custom Matlab script.

***In Vivo* Calcium Imaging**: Approximately 3 weeks after viral injection, fluorescence was checked using a custom modified UCLA miniscope (Cai et al., 2016) designed to hold a relay lens (1.8x 4.3 mm, Edmund Optics). A baseplate was then fixed over the implanted GRIN lens using dental cement. During behavioral testing, one reflective marker was adhered to top of the miniscope, and two were adhered to the side of the head bar to obtain pitch, roll and yaw angles as previously described. For calcium imaging analysis, all videos were preprocessed using Mosaic (Inscopix) for motion correction and spatial binning, and then subsequently analyzed using a custom MATLAB Script. Data were then processed using a constrained non-negative matrix factorization (CNMF) analysis) for denoising, deconvolving and demixing the data (Pnevmatikakis et al., 2016). This method subtracts out background fluorescence, and accurately localizes and segregates neuronal activity. Data were then imported into Neuroexplorer along with behavioral variables, where calcium activity and behavioral variables were compared. All further analyses were the same as electrophysiology.

**Histology**: To confirm viral expression and optic fiber and electrode placement, mice were transcardially perfused with 0.1M phosphate buffered saline (PBS) followed by 4% paraformaldehyde (PFA). To aid placement, heads were stored in 4% PFA with 30% sucrose for 72 hrs. Brains were then post-fixed for 24 hours in 30% sucrose prior to cryostat sectioning (Leica CM1850) at 60 µm coronally. Fiber, electrodes, and lens implantation sites were verified after sections were processed for the presence of cytochrome oxidase to visualize cytoarchitecture by rinsing in 0.1M PB before incubating in a diaminobenzidine, cytochrome C, and sucrose solution for ~2 hours at room temperature. Mounted cytochrome oxidase sections were then dehydrated in 200 proof ethanol, defatted in xylene, and coverslipped with cytoseal. To confirm eYFP and FusionRed coexpression in VTA *Vgat*+ cells, as well as labeling in the vicinity of dopaminergic cells of *Vgat-ires-Cre* transgenic mice, select sections were rinsed in 0.1M PBS for 20 min before being placed in a PBS-based blocking solution containing 5% goat serum and 0.25% Triton X-100 at room temperature for 1 hr. Sections were then incubated with a primary antibody (polyclonal rabbit anti-Vgat, 1:200 dilution, ThermoFisher, catalog no. PA5-27569; polyclonal rabbit anti-Dopamine transporter, 1:200 dilution, abcam, catalog no. 18441; polyclonal rabbit anti-Tyrosine hydroxylase, 1:200 dilution, Millipore, catalog no. AB152) in blocking solution overnight at 4 °C. Sections were then rinsed in PBS for 20 min before being placed in a secondary antibody used to visualize *Vgat, DAT,* or *TH* neurons in the VTA (goat anti-rabbit Alexa Fluor 594, 1:1000 dilution, abcam, catalog no. ab150080; goat anti-rabbit Alexa Fluor 488, 1:1000 dilution, abcam, catalog no. ab150077) for 1 hr at room temperature. Sections for fluorescent microscopy were mounted and immediately coverslipped with Fluoromount G with DAPI medium (Electron Microscopy Sciences; catalog no. 17984-24). Brightfield images for placement verification were acquired and stitched using an Axio Imager.M1 upright microscope (Zeiss) and fluorescent images were acquired and stitched using a Z10 inverted microscope (Zeiss) (Extended data Fig. 10).

**GABAergic Neuron Classification:** We used a Gaussian mixture model (GMM; Python sklearn package) to perform unsupervised clustering of neurons for classification without explicitly selecting a threshold, thus reducing bias. To perform this analysis, the mean waveforms were first normalized and fit with a principal component analysis. The top 10 components, which explained over 95% of the variance, were clustered using a GMM. Classifications were then used to analyze putative GABAergic neurons that were not optically tagged.

**Support Vector Regression Decoder:** Roll and yaw angle were decoded from the same horizontal tracking sessions, whereas pitch was decoded from a vertical tracking task. All of the data was binned into 50 ms intervals, and fit using support vector regression. As the number of neurons and the size of the training data affects decoding performance, we limited our analysis to sessions where we had at least 6 classified GABAergic neurons, and at least 3 ½ minutes of training data. For each of these sessions the model was trained on the first 60% of the data, and performance was evaluated on a contiguous set of held-out data (15%). Prior to fitting the model, the neural data was z-scored, and the head-angles were zero-centered. Using these methods, we were able to achieve good decoding performance, considering our relatively short sessions, and limited number of neurons compared to other decoding problems (Glaser et al., 2017).

**Statistical Analyses**: All statistical analyses were performed in MATLAB and GraphPad Prism. A power analysis was not conducted to determine sample size *a priori.*

*Correlation Analyses for Peri-event raster plots*: Data were aligned either towards the spout moving in the rightward or downward direction, and were binned in 50 ms windows and Gaussian smoothed with a filter width of 5 bins, within a total window of 5 s in Neuroexplorer using a peri-event raster. Output data were then sorted according to the minimum and maximum head angles and the corresponding neural activity and collapsed into 10 data points each. These data were then exported into GraphPad Prism, where a correlation analysis was performed. For population analyses, neural data and behavioral variables binned at 50 ms were normalized using a Z-score analysis performed using a custom MATLAB script. The normalized data from each animal were then averaged together, where a population average was obtained for both the behavioral variables and the neural data. For correlation analysis with calcium imaging data, the analysis was identical to the one described above, but data were binned at 100 ms and Gaussian smoothed with a filter width of 10 bins. To classify neurons as pitch, roll, or yaw, the highest correlational value between neural data and the relevant behavioral variable was chosen.

*Correlation Analyses Across Entire Behavioral Session*: For each animal, neural data and continuously monitored yaw, roll, or pitch behavioral variables for the entire recording session were constructed using 10 ms time bins in NeuroExplorer and exported to MATLAB. Behavioral variables and neural activity were then sorted in a pairwise fashion based on the behavioral maximum and minimum values. Forty separate bins were created for each sorted variable, then averaged, and a Pearson Correlation between them was performed. For selected neurons, neural activity and behavioral variable information were filtered to exclude data when the animal was not tracking the reward spout.

*Cross-Correlation Analyses:* For each animal, neural data and continuously monitored roll, yaw, or pitch behavioral variables for the entire recording session were extracted as described above. Using the behavioral variables and identified yaw, roll and pitch neurons from the correlation analyses, a custom MATLAB script was then utilized to perform cross-correlation analyses between the continuously monitored behavioral variables and neural activity. Cross-correlations were performed using MATLAB's *xcorr* function, using the behavioral variable as the reference time-series, and the neural data as the shifted time series. Latencies were determined by finding the lag to the maximum value of the cross-correlation for positively correlated neurons and the lag to the minimum value for anti-correlated neurons. Cross-correlations were Z-score normalized and averaged together across data sets from all animals to obtain population data.

**Code and Data availability**

All data and scripts for MATLAB are available upon request.

### Supplementary Figures

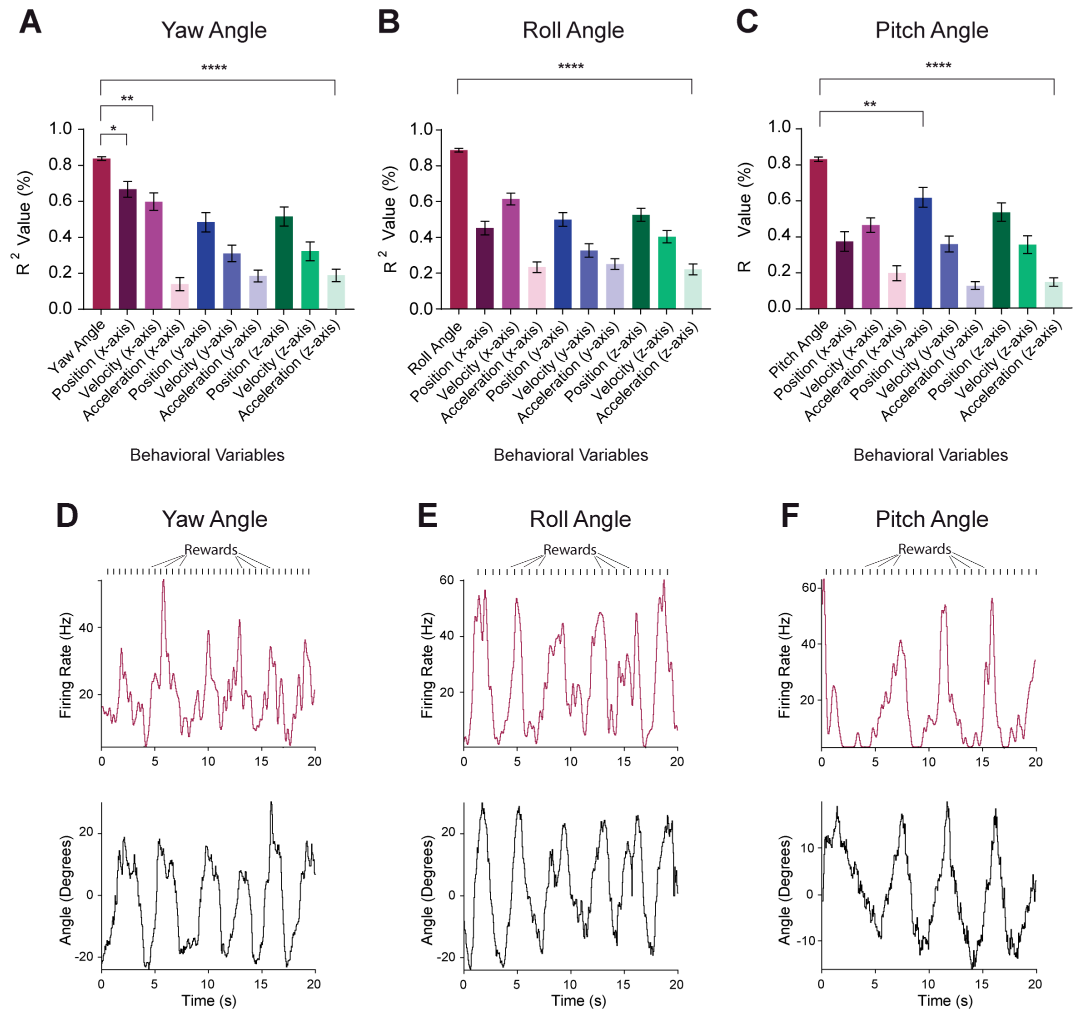


**Figure S1. Analyses for all other behavioral variables from pitch, roll and yaw neurons.**

**(A-C)** Correlation between other relevant behavioral variables and pitch, roll and yaw neurons showing a significant difference between all other behavioral variables.

**(A)** Neurons classified as Yaw angle neurons had a significantly higher correlational value than all other behavioral variables (one-way ANOVA, $p < 0.05$, $n = 26$).

**(B)** Roll Angle neurons had a significantly higher correlational value compared to other behavioral variables (one-way ANOVA, $p < 0.05$, $n = 62$).

**(C)** Pitch angle neurons had a significantly higher correlational value than all other behavioral variables (one-way ANOVA, $p < 0.05$, $n = 36$).

**(D-F)** Raw neural traces with their respective angle with corresponding reward times. Individual neurons did not show a clear relationship with reward.

**(D)** Raw traces for a representative yaw angle neuron and yaw angle.

**(E)** Raw traces for a representative roll angle neuron and roll angle

**(F)** Raw traces for a representative pitch neuron and pitch angle (* values reflect $p$ values adjusted for multiple comparisons using Dunnett's multiple comparison's test. * $p < 0.05$; ** $p < 0.01$; **** $p < 0.0001$).

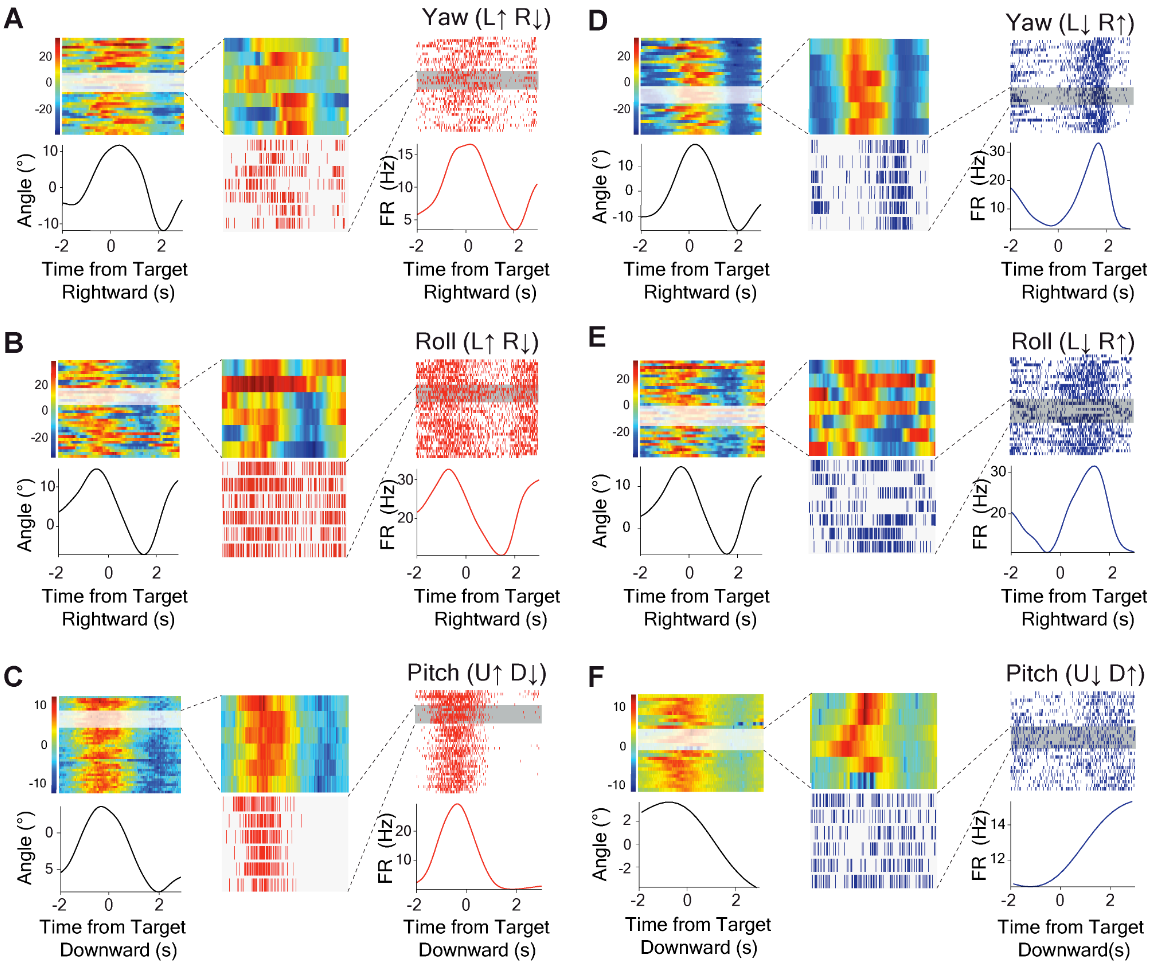


**Figure S2. Single-trial examples from representative neurons in Figure 1.**

Single trial examples demonstrating the robust correlation between neural activity and head angle**.** Head angle is on the left and VTA GABAergic neural activity is on the right. Select trials are shown in the middle.

**(A)** Single-trial perievent rasters and corresponding behavioral traces for representative Yaw (L↑ R↓) neuron (Seven trials; *PC*, $r^2 = 0.89$, $p < .0001$).

**(B)** Single-trial perievent rasters and corresponding behavioral traces for representative Roll (L↑ R↓) neuron (Six trials; *PC*, $r^2 = 0.99$, $p < .0001$).

**(C)** Single-trial perievent rasters and corresponding behavioral traces for representative Pitch (U↑ D↓) neuron (Six trials; *PC*, $r^2 = 0.93$, $p < .0001$).

**(D)** Single-trial perievent rasters and corresponding behavioral traces for (L↓ R↑) neuron (Six trials; *PC*, $r^2 = 0.80$, $p < .01$).

**(E)** Single-trial perievent rasters and corresponding behavioral traces for representative Roll (L↓ R↑) neuron (Seven trials; *PC*, $r^2 = 0.98$, $p < .0001$).

**(F)** Single-trial perievent rasters and corresponding behavioral traces for representative Pitch (U↓ D↑) neuron (Seven trials; *PC*, $r^2 = 0.91$, $p < .0001$).

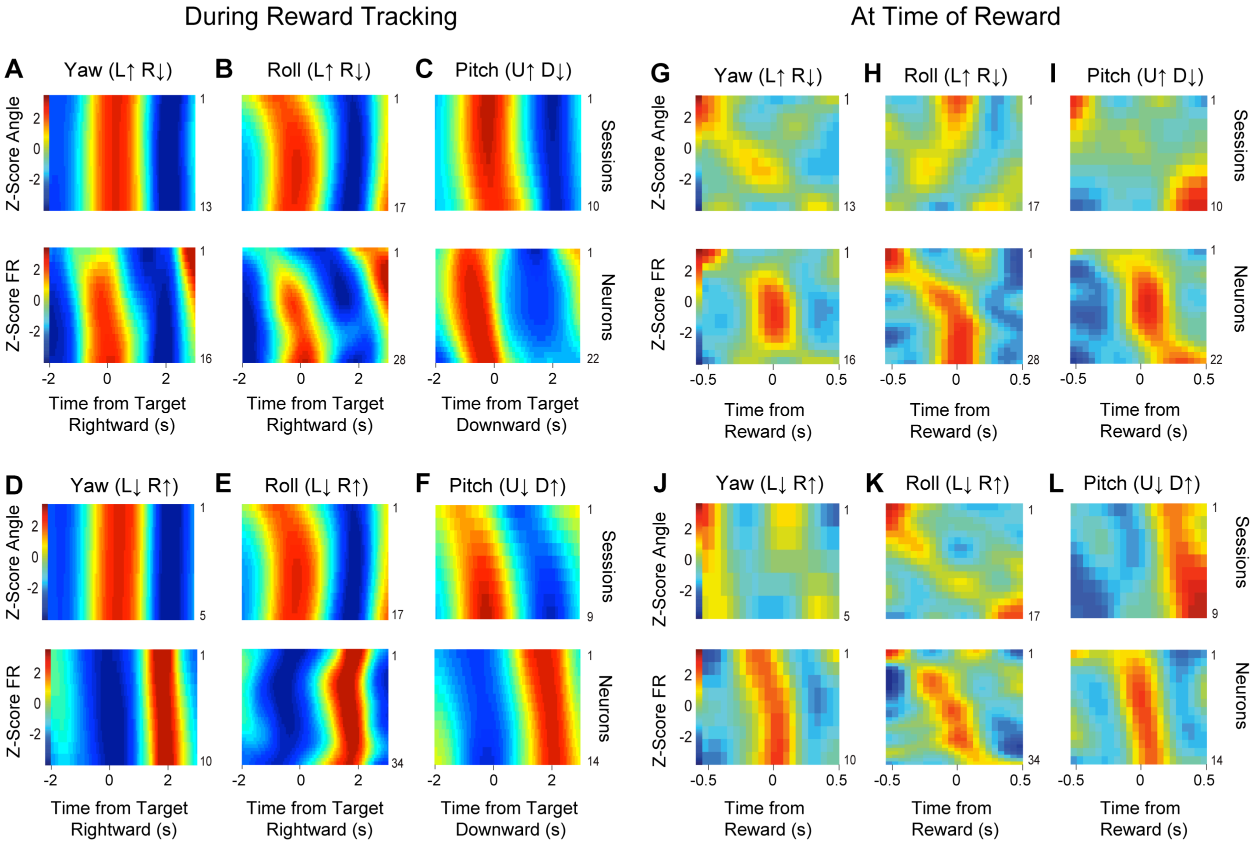


**Figure S3. Population heat maps of electrophysiological data for pitch, roll and yaw during reward tracking and at the time of reward consumption**.

**(A)** Yaw (L↑ R↓) angle population heat maps during tracking behavior. *Top*) Normalized yaw angle across 13 behavioral sessions. *Bottom*) Normalized firing rate of Yaw (L↑ R↓) neurons ($n$ = 16).

**(B)** Roll (L↑ R↓) angle population heat maps during tracking behavior. *Top*) Normalized roll angle across 17 behavioral sessions. *Bottom*) Normalized firing rate of Roll (L↑ R↓) neurons ($n$ = 28).

**(C)** Pitch (U↑ D↓) angle population heat maps during tracking behavior. *Top*) Normalized pitch angle across 10 behavioral sessions. *Bottom*) Normalized firing rate of Pitch (U↑ D↓) neurons ($n$ = 22).

**(D)** Yaw (L↓ R↑) angle population heat maps during tracking behavior. *Top*) Normalized yaw angle across 5 behavioral sessions. *Bottom*) Normalized firing rate of Yaw (L↓ R↑) neurons ($n$ = 10).

**(E)** Roll (L↓ R↑) angle population heat maps during tracking behavior. *Top*) Normalized roll angle across 17 behavioral sessions *Bottom*) Normalized firing rate of Roll (L↓ R↑) neurons (*n* = 34).

**(F)** Pitch (U↓ D↑) angle population heat maps during tracking behavior. *Top*) Normalized pitch angle across 9 behavioral sessions *Bottom*) Normalized firing rate of Pitch (U↓ D↑) neurons (*n* = 14).

**(G)** Yaw (L↑ R↓) angle population heat maps at time of reward. *Top*) Normalized yaw angle across 13 behavioral sessions. *Bottom*) Normalized firing rate of Yaw (L↑ R↓) neurons (*n* = 16).

**(H)** Roll (L↑ R↓) angle population heat maps at time of reward. *Top*) Normalized roll angle across 17 behavioral sessions. *Bottom*) Normalized firing rate of Roll (L↑ R↓) neurons (*n* = 28).

**(I)** Pitch (U↑ D↓) angle population heat maps at time of reward. *Top*) Normalized pitch angle across 10 behavioral sessions. *Bottom*) Normalized firing rate of Pitch (U↑ D↓) neurons (*n* = 22).

**(J)** Yaw (L↓ R↑) angle population heat maps at time of reward. *Top*) Normalized yaw angle across 5 behavioral sessions. *Bottom*) Normalized firing rate of Yaw (L↓ R↑) neurons (*n* = 10).

**(K)** Roll (L↓ R↑) angle population heat maps at time of reward. *Top*) Normalized roll angle across 17 behavioral sessions *Bottom*) Normalized firing rate of Roll (L↓ R↑) neurons (*n* = 34).

**(L)** Pitch (U↓ D↑) angle population heat maps. *Top*) Normalized pitch angle across 9 behavioral sessions *Bottom*) Normalized firing rate of Pitch (U↓ D↑) neurons (*n* = 14).

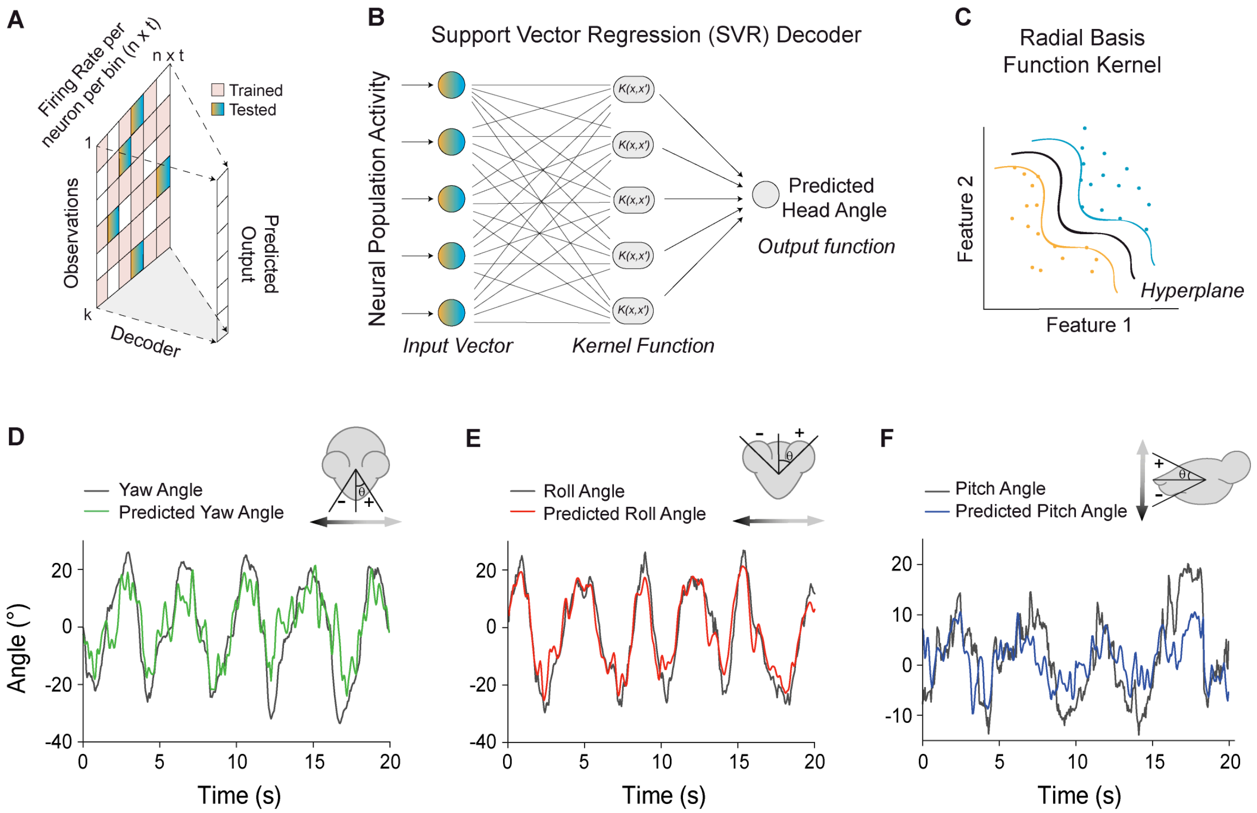


**Figure S4. Neural activity accurately predicts head angle using a Support Vector Regression (SVR) decoder**

**(A)** Schematic of a non-recurrent neural decoder used here to predict head-angle. It takes the firing rate of all neurons, *n*, across all time bins (50 ms), *t*. The first 60 % of the data was used to train the model, and a contiguous 15% of held-out data was then used to test the model.

**(B)** A Support Vector Regression (SVM) fits a kernel function to an input vector in order to predict the output vector, in this case the continuous head-angle measurements.

**(C)** A non-linear radial basis function kernel was used as the kernel function in the SVR decoder, which analyses patterns within the data set and separates them into subspaces by finding a hyperplane in the high-dimensional feature space.

**(D-F)** Schematic illustrated of the decoded head angle (*top*); Traces of head angle behavior during tracking, and the predicted head angle from the decoded neural activity (*bottom)*

**(D)** Representative model test on predicted yaw angle versus actual yaw angle from GABA population.

**(E)** Representative model test on predicted roll angle versus actual roll angle from GABA population.

**(F)** Representative model test on predicted yaw angle versus actual yaw angle from GABA population.

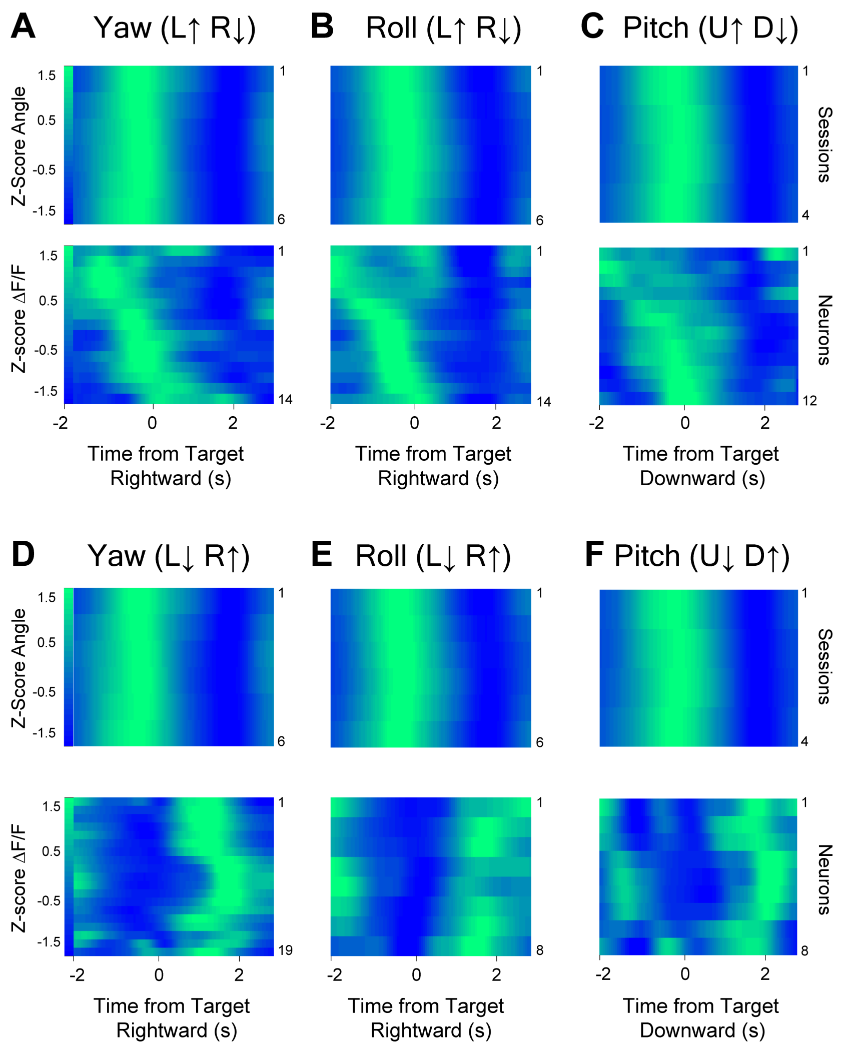


**Figure S5. Population heat maps of calcium imaging data for pitch, roll, and yaw.**

**(A)** Yaw (L↑ R↓) angle population heat maps. *Top*) Normalized yaw angle across 6 behavioral sessions. *Bottom*) Normalized ΔF/F of Yaw (L↑ R↓) neurons ($n$ = 14).

**(B)** Roll (L↑ R↓) angle population heatmaps. *Top*) Normalized roll angle across 16 behavioral sessions. *Bottom*) Normalized ΔF/F of Roll (L↑ R↓) neurons ($n$ = 14).

**(C)** Pitch (U↑ D↓) angle population heatmaps. *Top*) Normalized pitch angle across 4 behavioral sessions. *Bottom*) Normalized ΔF/F of Pitch (U↑ D↓) neurons ($n$ = 12).

**(D)** Yaw (L↓ R↑) angle population heat maps. *Top*) Normalized yaw angle across 6 behavioral sessions. *Bottom*) Normalized ΔF/F of Yaw (L↓ R↑) neurons ($n$ = 19).

**(E)** Roll (L↓ R↑) angle population heatmaps. *Top*) Normalized roll angle across 6 behavioral sessions. *Bottom*) Normalized ΔF/F of Roll (L↓ R↑) neurons ($n$ = 8).

**(F)** Pitch (U↓ D↑) angle population heatmaps. *Top*) Normalized pitch angle across 4 behavioral sessions. *Bottom*) Normalized ΔF/F of Pitch (U↓ D↑) neurons ($n$ = 8)

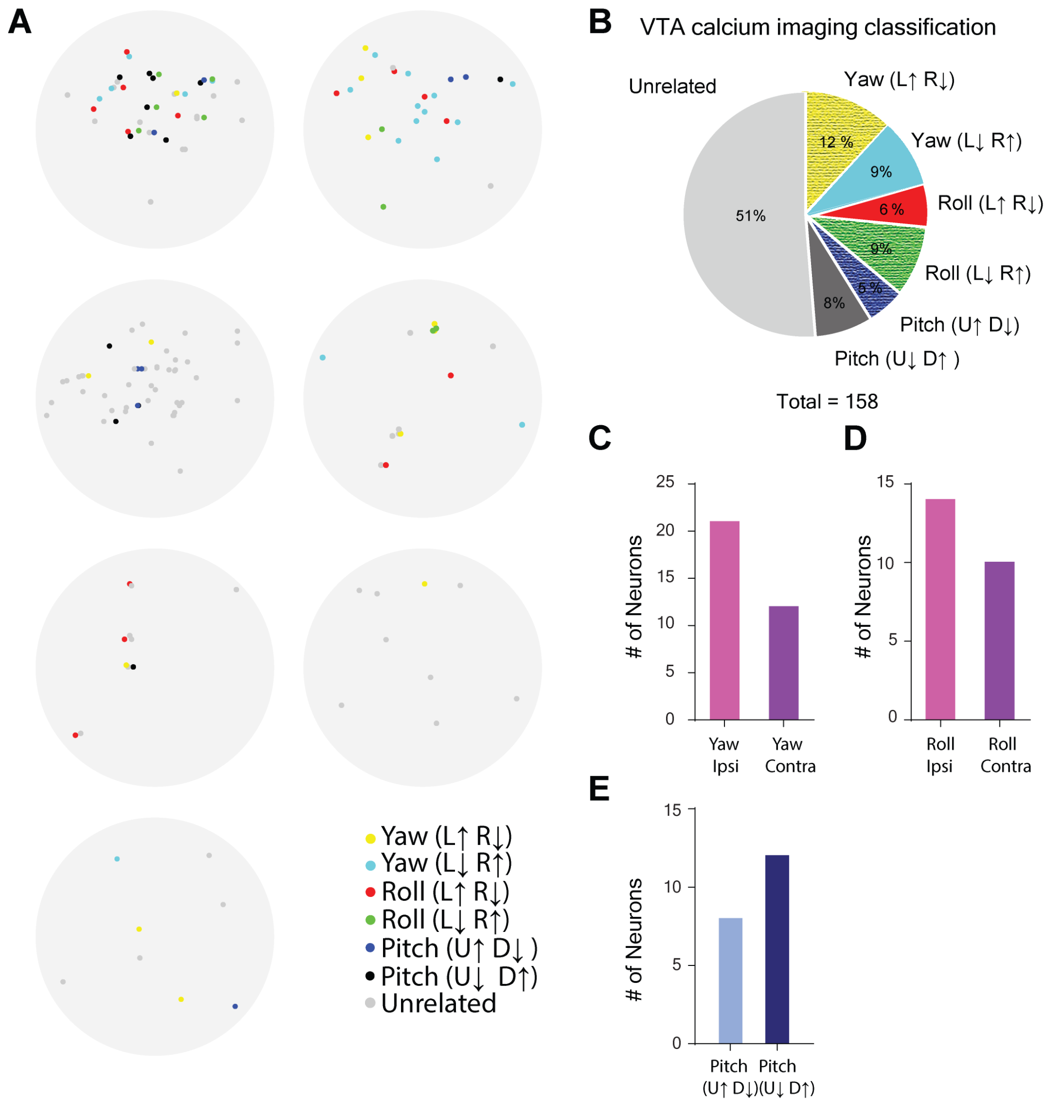


**Figure S6. Population summary of calcium imaging results**

**(A)** Topographical distribution of angle neurons within each animal from calcium imaging data ($n$ = 7).

**(B)** Breakdown of angle neuron percentages of calcium imaging neurons.

**(C)** Quantification of the number of Yaw angle neurons that increase their firing rate in the ipsiversive direction ($n$ = 21) or contraversive direction ($n$ = 12) in relation to the recording hemisphere.

**(D)** Quantification of the number of Roll angle neurons that increase their firing rate in the ipsiversive direction ($n$ = 14) or contraversive direction ($n$ = 10) in relation to the recording hemisphere.

**(E)** Quantification of the number of Pitch angle neurons that increase their firing rate in the upward direction ($n$ = 8) or downward direction ($n$ = 12).

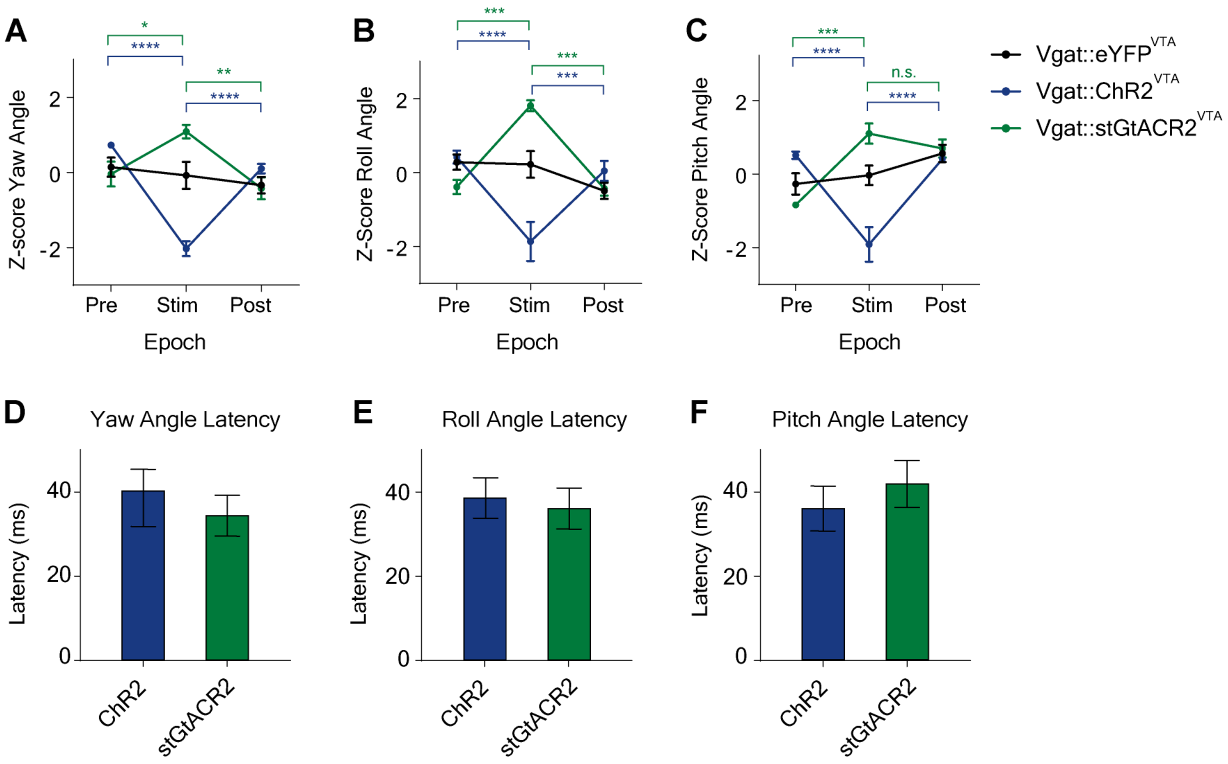


**Figure S7. Summary of optogenetic experiments**.

**(A)** Deviations of yaw angle over time during 500 ms of optogenetic excitation, inhibition and control. Yaw angle was significantly different during optogenetic excitation and inhibition than during pre and post period compared to control group (two-way RM ANOVA; Yaw Angle, Main effect of Group, $F_{(2,16)} = 12.16$, $p = 0.006$).

**(B)** Roll angle over time during 500 ms of optogenetic excitation, inhibition and control. Roll angle was significantly different during optogenetic excitation and inhibition than during pre and post period compared to control group (two-way RM ANOVA, Roll Angle, Main effect of Group, $F_{(2,16)} = 18.75$, $p < 0.0001$).

**(C)** Pitch angle over time during 500 ms of optogenetic excitation, inhibition and control. Pitch angle was significantly different during optogenetic excitation and inhibition than during pre and post period compared to control group (two-way RM ANOVA, Pitch Angle, Main effect of Group, $F_{(2,15)} = 12.20$, $p = 0.007$).

**(D)** Latency between optogenetic stimulation and deviation of yaw angle.

**(E)** Latency between optogenetic stimulation and deviation of roll angle.

**(F)** Latency between optogenetic stimulation and deviation of pitch angle. (* values reflect adjusted $p$ values from Tukey's *post-hoc* pairwise comparisons. * $p < 0.05$; ** $p = 0.004$; *** $p < 0.001$; **** $p < 0.0001$).

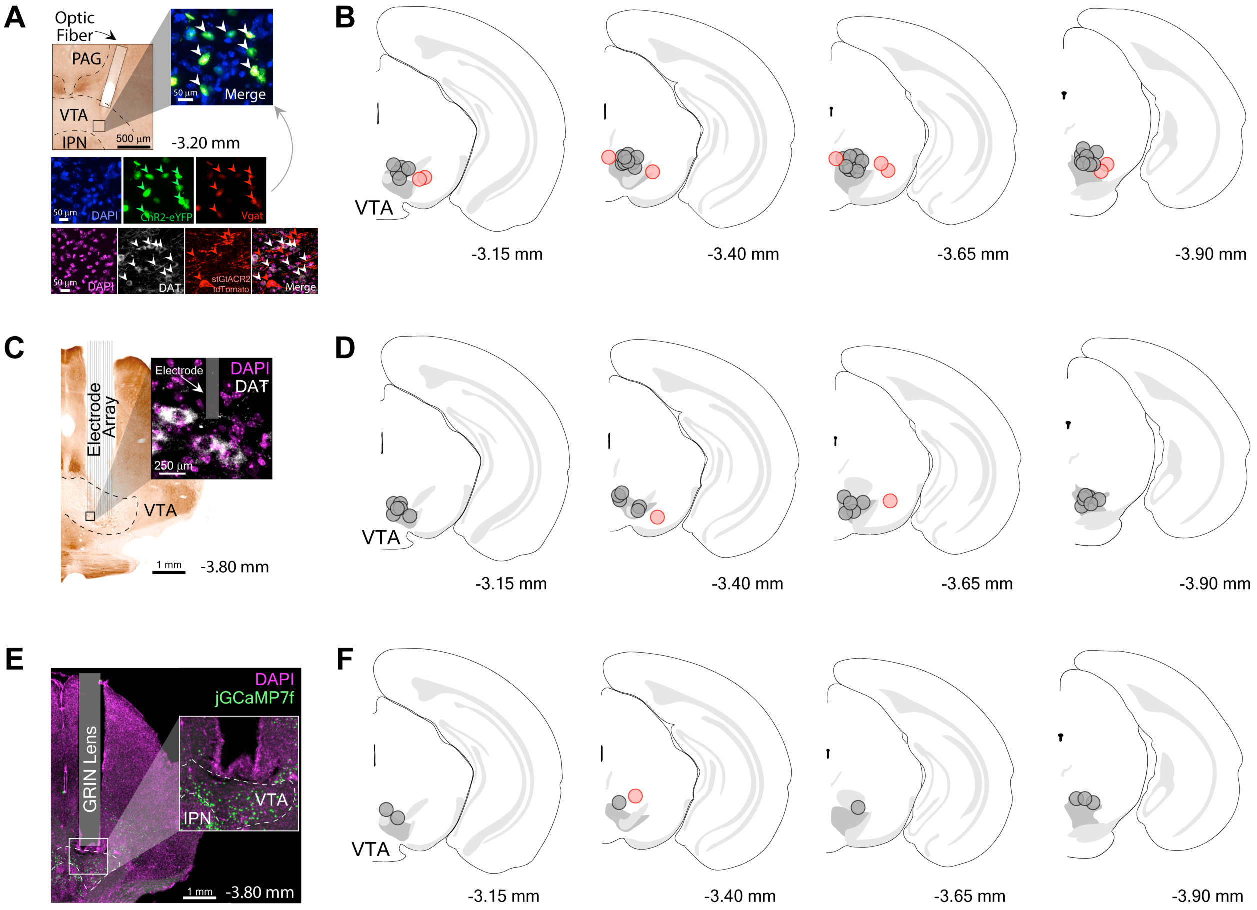


**Figure S8. Histological verification of VTA placement**

**(A)** Representative example of coronal section through the midbrain showing optic fiber placement above the VTA. Inset shows eYFP infected Vgat+ neurons in the VTA.

**(B)** Optic fiber placements for VTA optogenetic experiments ($n$ = 19).

**(C)** Representative example of coronal section through the midbrain showing electrode array placement into the VTA. Inset shows electrode tip co-localization with dopaminergic (DAT) neurons in the VTA.

**(D)** Electrode array placement into the VTA ($n$ = 22).

**(E)** GRIN lens placement above the VTA for calcium imaging experiments ($n$ = 7). Black circles denote data points included in study. Red circles denote excluded data points. Coordinates with respect to bregma shown beneath coronal schematics.

### Supplemental Videos

**Video S1.**
Overhead view of continuous horizontal tracking by a mouse wearing a wireless headstage, showing fluctuations in yaw and roll angles.

**Video  S2.**
Side view of continuous horizontal tracking by a mouse wearing a wireless headstage, showing fluctuations in yaw and roll angles.

**Video S3.**
Side view of continuous vertical tracking by a mouse wearing a wireless headstage, showing fluctuations in pitch angle.

**Video S4.**
Front and side views of effect of unilateral excitation of VTA GABAergic neurons in a freely moving *Vgat-ires-Cre* mouse that received DIO-ChR2 virus into the VTA. Videos show two different stimulation trials, aligned to laser onset time. Stimulation duration is 500 ms.

**Video S5.**
Front and side views of unilateral inhibition of VTA GABAergic neurons in a freely moving *Vgat-ires-Cre* mouse that received DIO-stGtACR2 virus into the VTA. Videos show two different stimulation trials, aligned to laser onset time. Stimulation duration is 200 ms.
